# Targeting Cellular Pseudo-Senescence to Overcome PARP inhibitors Resistance in *BRCA1*-Mutated Breast Cancer

**DOI:** 10.64898/2026.06.25.734617

**Authors:** Zhen Tian, Srinivas Chatla, Shreya Indulkar, Yang Liu, Alexander Morris, Bac Viet Le, Dongwook Kim, Xiaoyu Wei, Yun Liao, Dan Yang, Anthony Pompetti, Gennaro Calendo, Congli Wang, Tina B. Edmonston, Zhenkun Lou, Lu Chen, Tomasz Skorski, Liewei Wang, Jian Huang

## Abstract

Poly (ADP-ribose) polymerase (PARP) inhibitors (PARPi) are a mainstay therapy for homologous recombination (HR)-deficient cancers; however, resistance remains a major clinical challenge. Previously, through a genome wide CRISPR screen, we identified *ZNF251* haploinsufficiency as a novel driver of PARPi resistance. In *BRCA1*-mutant (*BRCA1*^mut^) cells, *ZNF251* deficiency led to HR hyperactivation, conferring PARPi resistance that could be reversed by RAD51 inhibition. In this study, we further show that *ZNF251* deficiency induces replication stress and a pseudo-senescence state, in which cells exhibit molecular and phenotypic markers of senescence while retaining proliferative capacity. Because senomorphic and senolytic therapies can target senescent cells, we tested whether these approaches could overcome PARPi resistance in *ZNF251*-deficient breast cancer cells. Critically, targeting this senescence-like state with either senomorphic agents, such as cytokine inhibitors, or senolytic agents, such as BCL-2 and BCL-XL inhibitors, overcame PARPi resistance *ex vivo* and *in vivo*. Importantly, pseudo-senescence was also observed in other PARPi-resistant contexts driven by HR hyperactivation, including *53BP1*- and *Shieldin*-mutant cells, suggesting that it may represent a broader mechanism underlying PARPi resistance in breast cancer. Furthermore, in two olaparib-resistant, *BRCA*^mut^ triple-negative breast cancer organoid models, treatment with DT2216, a BCL-XL–targeting PROTAC, sensitized both models to olaparib. Together, our work defines a novel pathway linking HR hyperactivation, replication stress, and pseudo-senescence, and positions both senomorphic and senolytic therapies as promising strategies to overcome PARPi resistance in *BRCA1*^mut^ breast cancer.

**Highlights:**

- *ZNF251* deficiency drives PARP inhibitor resistance through HR hyperactivation, replication stress, and pseudo-senescence in *BRCA1*-mutant breast cancer.
- Senomorphic and senolytic therapies overcome PARP inhibitor resistance *in vitro*, *in vivo*, and in patient-derived organoid models.
- Pseudo-senescence represents a shared vulnerability of HR-hyperactivated PARPi-resistant cancers and can be therapeutically targeted.

## Introduction

Poly(ADP-ribose) polymerase 1 (PARP1) plays a central role in DNA damage repair, and PARP inhibitors (PARPi) have become an important treatment for cancers with homologous recombination (HR) deficiency, particularly those harboring *BRCA1* or *BRCA2* mutations. By exploiting synthetic lethality, PARPi selectively target HR-deficient tumor cells and have demonstrated significant clinical benefit in *BRCA*-mutant breast and ovarian cancers[1, 2]. Beyond *BRCA* mutations, PARPi sensitivity has established the concept of “BRCAness” as a predictive biomarker across HR-deficient tumors[3].

Despite their clinical success, resistance to PARPi remains a major challenge. Approximately 40–70% of patients eventually develop resistance through diverse mechanisms, many of which overlap with resistance to platinum-based chemotherapy[4–6]. Identified resistance mechanisms include reduced intracellular drug availability, altered PARP activity, restoration or hyperactivation of HR repair, and stabilization of stalled replication forks. Loss of DNA repair factors such as *53BP1* and components of the Shieldin complex promotes PARPi resistance by restoring HR function[7, 8]. Recently, we identified *ZNF251* in an unbiased genome-wide CRISPR screen and demonstrated that heterozygous loss of *ZNF251* confers PARPi resistance in *BRCA1*-mutant breast and ovarian cancer models through HR hyperactivation[9]. However, the downstream consequences of HR overactivation and their contribution to PARPi resistance remain poorly understood.

Cellular senescence is a stress-induced state characterized by stable cell-cycle arrest and the secretion of pro-inflammatory factors collectively known as the senescence-associated secretory phenotype (SASP). While senescence can suppress tumor growth, SASP factors may promote tumor progression, therapeutic resistance, and remodeling of the tumor microenvironment[10–12]. A related state, termed pseudo-senescence, is characterized by the acquisition of senescence-associated features while retaining proliferative capacity and tumor-promoting potential[13–15]. Emerging evidence suggests that pseudo-senescent cells can survive anticancer therapies and support resistant tumor growth through SASP-mediated signaling[14, 15].

Targeting senescence-associated vulnerabilities has therefore emerged as a promising therapeutic strategy[16]. Senolytic agents, including BCL-2 and BCL-XL inhibitors[17, 18], selectively eliminate senescent cells by disrupting pro-survival pathways, whereas senomorphic agents suppress SASP-mediated signaling[13]. Although conventional BCL-XL inhibitors are limited by thrombocytopenia[19, 20], the BCL-XL–targeting PROTAC DT2216 was developed to selectively degrade BCL-XL while reducing platelet toxicity[21].

Although senescence has been implicated in therapy resistance, the role of pseudo-senescence in PARPi-resistant breast cancer remains largely unexplored. Whether HR hyperactivation drives replication stress and pseudo-senescence, and how SASP signaling contributes to PARPi resistance, has not been systematically investigated.

Here, we show that *ZNF251* deficiency induces replication stress and pseudo-senescence, leading to PARPi resistance in *BRCA1*-mutant breast cancer. We further demonstrate that targeting senescence-associated pathways with senomorphic or senolytic therapies restores PARPi sensitivity *in vitro* and *in vivo*. Notably, pseudo-senescence was also observed in *53BP1*- and *Shieldin*-deficient models, suggesting that it represents a broader mechanism of PARPi resistance. Furthermore, DT2216 restored olaparib sensitivity in two olaparib-resistant *BRCA*-mutant triple-negative breast cancer organoid models. Together, our findings establish a mechanistic link between HR hyperactivation, replication stress, and pseudo-senescence, and identify senescence-targeted therapies as a potential strategy to overcome PARPi resistance in *BRCA1*-mutant cancers.

## Materials and Methods

All key resources used in this study are listed in the Key Resources Table (Supplementary Table 1).

### Cell lines and cell culture

MDA-MB-436 and HCC1937 cells were purchased from ATCC and maintained in DMEM supplemented with 10% fetal bovine serum (FBS) and 1% penicillin/streptomycin. RPE1-hTERT *SHLD*-WT, *SHLD1*-KO, and *SHLD2*-KO cells were generously provided by Dr. Daniel Durocher (Lunenfeld-Tanenbaum Research Institute, Toronto, Canada). RPE1-hTERT cells were maintained in DMEM/F12 supplemented with 10% FBS and 1% penicillin/streptomycin. U2OS cells were maintained in McCoy’s 5A supplemented with 10% FBS and 1% penicillin/streptomycin. All cell lines used in this study were authenticated by short tandem repeat (STR) profiling (Labcorp, PowerPlex 16 HS) in October 2026 and compared with Cellosaurus reference profiles using CLASTR; all lines met the ≥80% criterion of ANSI/ATCC ASN-0002-2022 (Supplementary Table 2 and Supplementary materials). Cell morphology and relevant molecular characteristics were also routinely monitored during culture. All cell lines were tested for mycoplasma contamination and confirmed to be mycoplasma-free before use in the experiments.

### Mice and *in vivo* studies

Female NOD/SCID/IL-2Rγ (NSG) mice aged 6-8 weeks were obtained from The Jackson Laboratory (stock no. 005557; RRID:IMSR_JAX:005557). All animal experiments were approved by the Cooper University Healthcare Institutional Animal Care and Use Committee (IACUC; protocol No. 25-014). Mice were housed under pathogen-free conditions, with five animals per cage, at 26 °C, under a 12-h light/12-h dark cycle. Autoclaved food and water were provided *ad libitum*. Environmental enrichment consisted of nesting material and shelters. Animals were acclimatized for 15 days before experimental procedures.

Each mouse received subcutaneous injections of 1 × 10 MDA-MB-436 cells in 100 μL saline per flank, with WT and *ZNF251*-knockdown cells on opposite sides. Approximately four weeks later, when both tumors reached 20–30 mm³, mice were randomly allocated to vehicle, olaparib, navitoclax, or combination treatment (n = 4 mice/group; 16 mice total). Group sizes were based on our previous study demonstrating olaparib resistance in *ZNF251*-knockdown tumors[9]. No animals or data were excluded, and no attrition occurred.

Olaparib (10 mg/kg in 5% DMSO/saline) was administered intraperitoneally daily, and navitoclax (50 mg/kg in a 1:1 mixture of 30 mM HCl and 1% methylcellulose; final concentrations: 15 mM HCl and 0.5% methylcellulose) was administered by oral gavage every two days, both for four weeks. Each administration had a volume of 200 μL. Combination-treated mice received both schedules; controls received the corresponding vehicles at matched volumes, routes, and schedules. Doses were based on previous studies[22–24]. Tumor dimensions were measured weekly using calipers; volume was calculated as V = L × W²/2, where L and W are the longest and perpendicular diameters. The primary outcome was tumor growth; final tumor volume and weight were additional outcomes. Statistical methods accounting for paired tumors and repeated measurements are described in the Statistical Analysis section.

Tumor cells were injected without anesthesia. Animal health and tumors were monitored daily by the changes in body weight, activity, posture, grooming, food and water intake, and tumor ulceration. No analgesia or additional supportive care was administered. Humane endpoints included a combined tumor diameter >2 cm, body-weight loss of ≥20% from baseline, tumor ulceration, or persistent signs of distress. Animals meeting humane-endpoint criteria were to be euthanized immediately. No animals reached these criteria during the study. No adverse events or animal losses were observed during the study. After four weeks of treatment, mice were euthanized by CO₂ inhalation, followed by cervical dislocation. Tumors were excised, photographed, weighed, and processed for following analysis.

### Patient-derived organoids (PDXOs)

The organoid models used in this study were generated from previously described patient-derived xenograft (PDX) models[25]. We therefore refer to them as PDX-derived organoids (PDXOs) rather than patient-derived organoids. Human tumor material was obtained under a protocol approved by the Mayo Clinic Institutional Review Board (IRB #11-007860; MC-1137) and written informed consent for research use of the specimens was obtained from all participants.

TNBC PDX tumors were maintained in NSG mice until harvest. Fresh tumor tissue was dissociated with the Human Tumor Dissociation Kit (Miltenyi Biotec, #130-095-929) according to the manufacturer’s instructions and passed through a 70 μm cell strainer to generate a single-cell suspension. Residual mouse cells were depleted with the Mouse Cell Depletion Kit (Miltenyi Biotec, #130-104-694). The flow-through containing human tumor cells was collected, washed with PBS, and resuspended in DMEM supplemented with 10% FBS, 1% penicillin/streptomycin (Thermo Fisher Scientific, #15140-122), 1% GlutaMAX (Thermo Fisher Scientific, #35050-061), 1% MEM non-essential amino acids (Corning, #25-025-CI), and 1% sodium pyruvate (Corning, #25-000-CI). 10,000 Cells were seeded into 96-well PhenoPlates (PerkinElmer, #6055302) in an enriched organoid media, and PDXOs were allowed to establish for 1–3 days before treatment. Drug concentrations were chosen based on preliminary experiments showing an IC50 of approximately 5 μM. Organoid media: DMEM high glucose -Invitrogen – 11965-092 w/ phenol red was supplanted with 10%FBS, 1% pen/strep (Lifetech 15140-122), 1% glutamax (Lifetech 35050-061), 1% MEM NEAA (MT-25-025-CIRF), 1% sodium pyruvate (MT-25-000-CIRF)

DT2216 (Cat. No.: HY-130604) and Olaparib (Cat. No.: HY-10162) was obtained from MCE and used for organoid experiments. All drug experiments had 3 biological replicates and 4 technical repeats per concentration tested. Treatment was done for 72 hours and % Viability was standardized to DMSO control.

Treatment efficacy was assessed with the ATP-based CellTiter-Glo 3D Cell Viability Assay (Promega, #G9683) according to the manufacturer’s protocol. Plate reader used is Tecan Spark High performance Multi-Mode Microplate Reader. Before use, the PDXO models were confirmed to retain the *BRCA*-mutant, homologous recombination-deficient phenotype of the parental tumor by immunohistochemical (IHC) staining for γ-H2AX, RAD51, and Ki-67.

### Immunohistochemistry (IHC) staining of PDXOs

Paraffin-embedded PDX tumors collected from mice were used for immunohistochemistry (IHC). Slides were heated at 60 °C for 1 h, deparaffinized in Histon-clear, and rehydrated through graded ethanol (100%, 90%, 85%, and 70%) to ddH₂O. Antigen retrieval was performed in citrate buffer (Thermo Fisher Scientific, Waltham, MA, USA) at 95–98 °C for 20 min. After cooling and washing with TBS, endogenous peroxidase activity was quenched with 3% H₂O₂ (Sigma-Aldrich, St. Louis, MO, USA) for 10 min. Sections were blocked for 1 h and incubated with primary antibodies overnight at 4 °C, followed by secondary antibodies for 1 h. After TBST washing, sections were treated with an Avidin/Biotin Blocking Kit (Vector Laboratories, Burlingame, CA, USA) for 30 min and developed with DAB substrate (Cell Signaling Technology, Danvers, USA) for 1 min. Sections were counterstained with hematoxylin (Sigma-Aldrich) for 30 s, rinsed in tap water for 10 min, and mounted with UltraMount aqueous permanent mounting medium (Bio-Rad Laboratories, Hercules, CA, USA). Images were acquired using a ZEISS AXIOSCAN 7 (Carl Zeiss Microscopy GmbH, Jena, Germany) with 20× objectives.

### Inhibitors and cytotoxic drugs

Olaparib (Catalog# HY-10162), Navitoclax (Catalog# HY-10087), A-1331852 (Catalog# HY-19741), NF-κB inhibitor (BAY11-708242) (Catalog# HY-13453), Y-320 (IL-17A inhibitor) (Catalog# HY-15898), and LMT-28 (IL-6 receptor antagonist) (Catalog# HY-102084) were purchased from MedChem Express (MCE). All compounds were dissolved, aliquoted, and stored in accordance with the manufacturer’s instructions.

### Cell viability assay

For 2D viability experiments, cells were seeded at 1 × 10^4 cells/well in 100 μL of complete medium in 96-well plates and treated with the indicated compounds. Viability was assessed at 72 hr after treatment. Trypan blue dye exclusion was used for measuring cell viability, with viable cells quantified using manual hemocytometer. Growth curves were generated by measuring viable cell numbers at different doses of drugs and plotting cell number relative to the starting population or day 0, as appropriate. Both MTS (Promega, #G3582) and CCK-8 (APExBIO, #K1018) assays were used to further conform the cell viability results, following the manufacturers’ protocols; absorbance was measured at 490 nm and 450 nm respectively. For 3D PDXO experiments, viability was measured using CellTiter-Glo 3D, as described in the PDX-derived organoid section, rather than conventional colorimetric MTS/CCK-8 assays.

### Clonogenic survival assay

For the clonogenic assay, 10^4^ cells/well were seeded in a 48-well plate. After 24 hours, the culture medium was replaced with fresh medium containing individual or combined inhibitors, and cells were incubated for 4 days. Following treatment, cells were trypsinized and re-seeded at 8x10^3^ cells/well into a 6-well plate for colony formation. After 7 days, the medium was aspirated, and cells were washed with 1X PBS prior to fixation in 4% formaldehyde (ThermoFisher, F79-500) for 20 minutes. The fixative was removed, and cells were rinsed twice with Milli-Q water and stained with 0.5% (w/v) crystal violet (Sigma, C0775-25G) for 20 minutes. Excessive stains were rinsed off under running tap water. Images of the colonies were acquired, and colony counts were analyzed using ImageJ.

### Immunofluorescence (IF) and confocal microscopy

MDA-MB-436 WT and *ZNF251*KD3 cells were grown on coverslips, treated with olaparib at 5 μM for 72 h, and processed for immunofluorescence. Cells were fixed with 4% (v/v) paraformaldehyde for 20 min at 4°C, washed with PBS, and permeabilized with 0.5% (v/v) Triton X-100 for 10 min. Cells were blocked with PBS containing 3% BSA for 1 h at room temperature and incubated overnight at 4°C with phospho-Histone H2A.X (Ser139) (γH2AX) monoclonal antibody (Cell Signaling Technology, #9718; 1:200 working dilution) diluted in PBS containing 5% BSA. The next day, cells were washed three times with PBST and incubated for 1 h at room temperature with anti-rabbit Alexa Fluor 488 (Invitrogen, #A-11008; 1:1000 working dilution) secondary antibody. Cells were then washed three times with PBST and mounted using 20 μL of DAPI-containing mounting medium (Vectashield, #H-1200-10). Images were acquired using a Leica SP8 confocal microscope with a 63× oil-immersion objective under identical acquisition settings within each experiment and analyzed using ImageJ. For quantification, the integrated density of γH2AX staining was measured from three independent experiments using ImageJ software.

### DNA fiber assay for measuring replication stress

DNA fiber assay was performed as described previously with modifications[26]. Briefly, cells were treated with or without olaparib for 48 h, followed by stepwise incubation with 50 μM CldU (40 min), 4 mM HU (4 h), and 250 μM IdU (40 min). Cells were washed 3 times with PBS between each incubation step. Then, cells were harvested and resuspended in 1X PBS at a concentration of 1000 cells/μl. 2 μL of cell solution was lysed on slides (Superfrost Plus microscope slides, Fisher Scientific, cat# 12-550-15) with 200 mM Tris–HCl, 50 mM EDTA, 0.5% SDS, pH 7.4 buffer for 8 min. Slides were tilted to a 15-degree angle to spread fibers, followed by air drying and fixing in 3:1 methanol:acetic acid for 10 min, and 1 wash with water, followed by air drying. Then, slides were stored at −20° C until further use. Next, denaturation of DNA fibers was performed in 2.5 M HCl for 2.5 h, followed by 2X washing with PBS for 5 min before and after denaturation. Slides were then incubated with BSA solution (2% in PBS) for 40 min at room temperature before incubation with primary antibodies recognizing CldU (Abcam, cat# ab6326, dilution 1:300) and IdU (BD Biosciences, cat# 347580, dilution 1:100) for 2.5 h at room temperature, 3 times washed, then incubated with Alexa Fluor 488 and Alexa Fluor 594 conjugated secondary antibodies (ThermoFisher Scientific, cat# A11062 and A21470, respectively, dilutions 1:300) for 1 h at room temperature. Fibers were mounted after 3 washes with PBS. Images were acquired with a Leica SP8 LSM Confocal microscope, and fiber lengths were measured using ImageJ software. A minimum of 100 replication forks were measured per group, and DNA fiber lengths are presented as IdU/CldU length ratio.

### DNA damage/repair assays

HR was measured using the DR-GFP reporter cassette as described previously[27]. Briefly, the DR-GFP plasmid (pDRGFP plasmid, Addgene, #26475) was restriction digested with I-SceI restriction enzyme (New England Biolabs, #R0694S) for an hour at 37°C. Then, the digested pDRGFP plasmid and undigested pDsRed plamid were transfected with lipofectamine 2000 transfection reagent (Invitrogen, #11668027) into MDA-MB-436 WT, ZNF251KD and MDA-MB-436 cells reconstituted with the WT *BRCA1* gene. The cells were treated with DMSO or Olaparib (5 μM for 72 h) after the 24 h of transfection. Cells were analyzed 72 h later by flow cytometry to assess DSB repair activity. HR efficiency was calculated as the number of GFP-positive cells divided by the number of transfected DsRed-positive cells. Three independent experiments were performed, and the results were analyzed using GraphPad Prism software.

### Senescence-associated β-galactosidase (SA-β-gal) detection

The SA-β-gal activity was measured using the senescence assay kit (Abcam, #ab228562) and senescence beta-galactosidase staining Kit (Cell Signaling Technology, #9860S) following the manufacturer’s instructions. Briefly, cells were seeded in 6-well plates or 8-well chamber slides and treated with either vehicle control or olaparib for 72h. For flow cytometry-based analysis, cells were harvested after treatment, washed with PBS, and incubated with the provided fluorogenic substrate at 37°C for 1 h in the dark. Cells were then washed, resuspended in assay buffer, and analyzed by flow cytometry using the FITC channel. Data were analyzed using FlowJo software. For SA-β-gal staining, cells were washed once with PBS and fixed with 1× fixative solution for 15 minutes at room temperature. After fixation, cells were rinsed twice with PBS and incubated with freshly prepared β-galactosidase staining solution containing X-gal (final pH adjusted to 6.0) at 37°C overnight in a dry incubator without CO_2_. Senescent cells were identified by the presence of blue staining and visualized using a microscope at 200X magnification.

### *ZNF251* complementation experiment

Exponentially growing MDA-MB-436 *ZNF251* WT and KD cells were seeded in six-well plates (1 million cells/well) and transfected with pcDNA3.1 vector or human *ZNF251* on pcDNA3.1 plasmid carrying the neomycin resistance (neo) gene. After transfection with 1μg plasmids, the cells were selected with G418 (400 µg/ml) in the culture medium for 2 weeks to maintain the selection of neomycin-resistant cells to generate stably transfected cell lines. Cells were plated in 96-well plates at a density of 1× 10^4^ cells/well in triplicate. The next day, the transfected cells were treated with DMSO or olaparib for three days, and SA-β-gal assay was performed.

### Cell cycle analysis by flow cytometry

Cells were seeded in 6-well plates and treated with vehicle or olaparib for 72 h after seeding. Cells were then harvested and fixed in 70% ethanol at -20°C for 24 h. Following fixation, cells were washed with PBS to remove residual ethanol and incubated with Hoechst 33342 and Pyronin Y to assess DNA and RNA content, respectively. Cell cycle distribution was analyzed by flow cytometry, and cells were categorized into G0/G1, S, and G2/M phases based on combined DNA and RNA content. Data were analyzed using FlowJo software.

### Immunofluorescence analysis of γH2AX foci

Cells were seeded in 8-well chamber slides and exposed to 4 Gy X-ray irradiation using a Rad Source RS-2000 irradiator. Following irradiation, cells were allowed to recover for 0.5, 1, 3, 5, or 7 h and were then fixed with 4% paraformaldehyde for 20 min and permeabilized with 0.05% Triton-X 100 in PBS for 10 min. Then, cells were blocked with PBS containing 3% BSA for 1 h at room temperature and incubated overnight at 4°C with phospho-Histone H2A.X (Ser139) (γH2AX) monoclonal antibody (Cell Signaling Technology, #9718; 1:200 working dilution) diluted in PBS containing 5% BSA. The next day, cells were washed three times with PBST and incubated for 1 h at room temperature with anti-rabbit Alexa Fluor 488 (Invitrogen, #A-11008; 1:1000 working dilution) secondary antibody. Cells were then washed three times with PBST and mounted using 20 μL of DAPI-containing mounting medium (Vectashield, #H-1200-10). Images were acquired using a Leica SP8 confocal microscope with a 63× oil-immersion objective under identical acquisition settings within each experiment and analyzed using ImageJ. For quantification, the integrated density of γH2AX staining was measured from three independent experiments using ImageJ software.

### Cytokine array analysis

Cells were seeded in 6-well plates and treated with vehicle or olaparib for 24 h after seeding. Following treatment, culture-conditioned media were collected and centrifuged at 300-500 g for 5 min to remove cellular debris, and either used immediately or stored at -80°C until analysis. Secreted cytokine levels were measured using a cytokine array (RayBiotech, #AAH-CYT-5-4) according to the manufacturer’s instructions. Chemiluminescent signals were detected using the iBright 1500 imaging system and quantified with iBright analysis software.

### Real-time PCR (qPCR)

Total RNA was extracted from harvested cells using TRIzol reagent (Invitrogen, #15596026). RNA concentration and purity were assessed using a NanoDrop instrument (A260/A280). cDNA was synthesized from 1 μg of total RNA using the SuperScript IV VILO Master Mix (Invitrogen, #11756050) in a final reaction volume of 20 μL. Quantitative PCR was performed using Power SYBR Green Master Mix (Applied Biosystems, #4367659) on a QuantStudio 6 Flex Real-Time PCR System (Applied Biosystems). Primer sequences for IL-1α, IL-1β, IL-6, IL-8, TNF-α, p16INK4a, p21CIP1/WAF1, and the reference genes are provided in the List of Primers. Thermal cycling conditions were 95 °C for 2 min, followed by 40 cycles of 95 °C for 15 s and 60 °C for 1 min, followed by melt-curve analysis. Primer specificity was verified by melt-curve analysis and agarose gel electrophoresis. Amplification efficiencies for each target and reference assay were determined from serial cDNA dilution curves (E = 10^(−1/slope) − 1). Relative expression was calculated using the 2^(−ΔΔCt) method for assays meeting the efficiency criteria. Primary analyses were normalized to GAPDH, and results were confirmed using the geometric mean of *TBP*, *HPRT1*, and *PPIA* (Supplementary Fig. 3C). Because *GAPDH* expression can vary under metabolic and senescence-associated conditions, this independent normalization was used to exclude reference-gene artifacts. Three independent experiments were performed, each with three technical replicates per gene.

### Western blotting

Cells were directly harvested and lysed in RIPA buffer supplemented with 1× loading buffer. Protein samples were separated by SDS–PAGE and transferred onto nitrocellulose (NC) membranes. Membranes were blocked with 5% non-fat dry milk in TBST and subsequently incubated overnight at 4°C with primary antibodies against p16, p21, and Lamin B1 (Senescence Marker Antibody Sampler Kit #56062), p53 (#2527), and GAPDH (#2118), all purchased from Cell Signaling Technology (Danvers, MA, USA). Primary antibodies were diluted according to the manufacturer’s instructions. Following TBST washes, membranes were incubated with HRP-conjugated anti-rabbit secondary antibodies, and protein bands were detected using enhanced chemiluminescence. GAPDH served as the internal loading control. To ensure comparable protein loading across experimental groups without pre-quantification, initial lysates were analyzed, and sample loading volumes were empirically adjusted in subsequent run.

### CUT&RUN and chromatin analysis

CUT&RUN for H3K27me3 and H3K4me3 was performed on MDA-MB-436 ZNF251WT and ZNF251KD3 cells in two independent datasets (RepA and RepB), each with a matched IgG control. For ZNF251, CUT&RUN was attempted with two commercial antibodies (Proteintech 25601-1-AP; Invitrogen PA5-53774); neither gave specific signal enriched over IgG. Library prep was done using the NEB Ultra II kit (NEB E7645L) with single index barcodes (NEB E6609S). Libraries were sequenced paired-end and aligned to GRCh38RPGC using Bowtie2. Bigwigs and heatmap plots were generated using Deeptools. Peaks were called using MACS3 and analyzed in R using DiffBind.. Coverage tracks were generated as counts per million (CPM). Broad peaks were called for each sample against its matched IgG with MACS3 (--broad --broad-cutoff 0.1 -q 0.05 --max-gap 5000 --min-length 1000), and calls from all samples were merged within 5 kb to give region-level sets. For the domain-level analysis, blocks of at least 100 kb were merged across gaps of up to 1 Mb (411 domains). Domains were called gained or lost when the mean CPM changed by at least 0.1 (KD minus WT). In Fig. 8A, the log2 KD/WT ratio was centered on its median to correct for the global shift in signal, and regions at least 2-fold above or below the median were classed as gained or lost. Heatmaps and profiles were generated with deepTools, with each region scaled to equal length and 20 kb flanks. ENCODE ZNF251 ChIP-seq from HepG2 cells (ENCSR605KVV) is shown for reference only. Our CUT&RUN data and a side-by-side comparison with the ENCODE ZNF251 ChIP-seq will be deposited in GEO [accession to be added].

### Quantification and statistical analysis

Statistical analyses were performed using GraphPad Prism software. Unless otherwise stated, data are presented as mean ± SD from at least three independent biological experiments; technical replicates were averaged within each biological experiment and were not treated as independent observations. The biological replicate (n) and the statistical test used are reported for each experiment in the corresponding figure legend. For datasets with a sample size sufficient for distributional assessment, normality was evaluated using the Shapiro-Wilk test and graphical inspection of the data/residuals. Because formal normality tests have limited power at very small sample sizes (e.g., n = 3-4), statistical tests for such experiments were selected using non-parametric methods (Mann-Whitney), which do not assume a normal distribution. Homogeneity of variance was assessed using Brown-Forsythe test. Two-group comparisons meeting parametric assumptions were analyzed using a two-tailed unpaired Student’s t test; when variances were unequal, Welch’s t test was used. Two-group comparisons that did not reasonably satisfy parametric assumptions were analyzed using Mann-Whitney test. For comparisons involving more than two groups, one-way ANOVA with Tukey’s post hoc multiple-comparison procedure was used when parametric assumptions were met; Welch ANOVA or the Kruskal-Wallis test with Dunn’s post hoc test was used when variance or distributional assumptions were not met. For experiments involving more than one independent factor (for example, genotype and treatment), a two-way ANOVA followed by Šídák’s (or Tukey’s) multiple comparisons test was applied where appropriate All tests were two-sided unless otherwise specified. Exact P values are reported when feasible; P < 0.05 was considered statistically significant. Significance symbols are defined as *P < 0.05, **P < 0.01, ***P < 0.001, and ****P < 0.0001. No data points or animals were excluded from any analysis, and no missing data were imputed.

## Results

### *ZNF251*KD caused cellular pseudo-senescence and led to PARPi resistance in *BRCA1*^mut^ breast cancer

Previously, through an unbiased genome-wide CRISPR screen, we identified *ZNF251* as a gene whose deficiency confers strong resistance to PARPis in MDA-MB-436 cells, a *BRCA1*-deficient triple-negative breast cancer (TNBC) cell line. Importantly, ZNF251’s functional role extends beyond PARPi resistance. We found that *ZNF251*KD also confers strong resistance to platinum-based chemotherapy, particularly cisplatin[9]. In MDA-MB-436 cells, the cisplatin IC50 was 15.38-28.88 μM across three independent *ZNF251*KD clones, compared with 6.3 μM for *ZNF251*WT cells. Mechanistic studies showed that *ZNF251*KD induces HR hyperactivation, thereby driving PARPi resistance in *BRCA1*^mut^ breast cancer[9]. These findings establish ZNF251 as an important novel regulator of HR-mediated PARPi sensitivity in *BRCA1*^mut^ breast cancer. To further elucidate the molecular mechanisms underlying *ZNF251*KD–mediated PARPi resistance, we performed RNA-seq analysis of olaparib-treated *ZNF251*WT and KD *BRCA1*^mut^ breast cancer cells compared with controls. Differential gene expression analysis showed distinct transcriptional responses to olaparib treatment between *ZNF251*WT and *ZNF251*KD cells. Compared with *ZNF251*WT cells, *ZNF251*KD cells exhibited a markedly attenuated transcriptional response to olaparib, consistent with their olaparib-resistant phenotype (Fig. 1A). Interestingly, Gene Set Enrichment Analysis (GSEA) of the SAUL_SEN_MAYO senescence signature showed significant enrichment of senescence-associated genes in vehicle-treated *ZNF251*KD cells compared with WT cells (NES=2.12, p=1×10⁻¹□) (Fig. 1B), indicating that *ZNF251* deficiency promotes a basal senescence-like transcriptional program. Consistent with PARPi-induced stress responses[28, 29], olaparib treatment enriched the senescence signature in WT cells (NES=1.45, p=0.0183). In contrast, olaparib-treated *ZNF251*KD cells exhibited significant negative enrichment relative to both vehicle-treated *ZNF251*KD cells (NES=−2.28, p=1×10⁻¹□) and olaparib-treated WT cells (NES=−1.75, p=0.00024) (Fig. 1B), suggesting an altered or attenuated senescence-associated transcriptional response. Together, these findings support a model in which *ZNF251* deficiency promotes a pre-existing pseudo-senescent state that may facilitate adaptation to PARPi–induced replication stress and contribute to PARPi resistance (Supplementary Fig. 1). In addition, Gene Set Variation Analysis (GSVA) pathway analysis showed enrichment of multiple senescence-related pathways, including oncogene-induced senescence, TP53 signaling, oxidative stress-induced senescence, and senescence-associated metabolic programs in *ZNF251*KD cells (Supplementary Fig. 2). Together, these findings suggest that *ZNF251*KD induces a senescence-associated transcriptional reprogramming state that may contribute to adaptive resistance to PARP inhibition.

**Figure 1.**
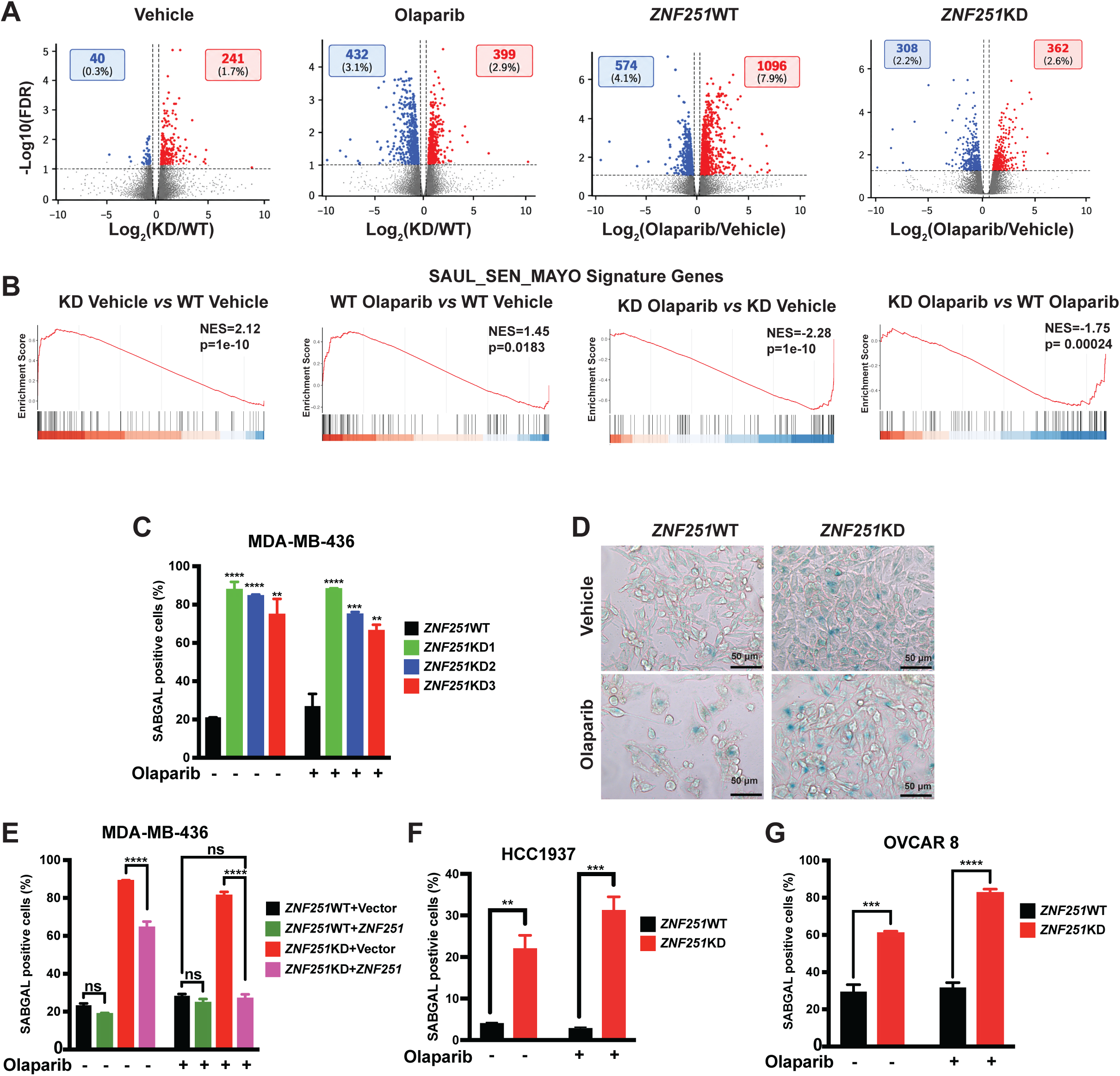
*ZNF251*KD induces senescence-associated transcriptional signatures and cellular senescence in *BRCA1*^mut^ breast cancer cells. (A) Volcano plot showing differential gene expression associated with *ZNF251* deficiency and olaparib treatment. (B) Gene set enrichment analysis (GSEA) of the SAUL_SEN_MAYO senescence signature in *ZNF251*WT and *ZNF251*KD cells treated with vehicle or olaparib. (C) Quantification of SA-β-gal-positive breast cancer cells in *ZNF251*WT and three independent *ZNF251*KD clones by flow cytometry (%). (D) Representative SA-β-gal staining images of *ZNF251*WT and *ZNF251*KD breast cancer cells treated with DMSO or olaparib. Scale bars, 50 μm. (E) Ectopic expression of wild-type *ZNF251* cDNA in *ZNF251*WT or *ZNF251*KD cells significantly reduced the percentage of SA-β-gal-positive cells. (F) Quantification of SA-β-gal-positive HCC1937 breast cancer cells in *ZNF251*WT and *ZNF251*KD groups by flow cytometry (%). (G) Quantification of SA-β-gal-positive OVCAR8 ovarian cancer cells in *ZNF251*WT and *ZNF251*KD groups by flow cytometry (%). Data represent the means ± SD of three replicates.

Consistent with the enrichment of senescence-associated gene signatures, three independent *ZNF251*KD clones of *BRCA1*^mut^ breast cancer cells showed a significant increase in the proportion of SA-β-gal–positive cells compared with *ZNF251*WT cells. These cells also exhibited pronounced morphological changes, including enlarged cell size and a flattened morphology (Fig. 1C and D). Importantly, ectopic expression of wild-type *ZNF251* cDNA in *ZNF251*KD cells significantly reduced the percentage of SA-β-gal–positive cells, confirming that *ZNF251* deficiency contributes to the senescence-associated phenotype (Fig. 1E). To determine whether this phenotype extends beyond MDA-MB-436 cells, we examined additional *BRCA1*-deficient cancer models. *ZNF251* knockdown significantly increased SA-β-gal activity in HCC1937 cells, an independent *BRCA1*-mutant breast cancer cell line (Fig. 1F), and in OVCAR8 cells, an ovarian cancer cell line with *BRCA1* promoter methylation and reduced *BRCA1* expression (Fig. 1G). We next asked which features of the canonical senescence program *ZNF251*KD cells acquire and which they lack. Persistent DNA damage is a major trigger of senescence[30], and *ZNF251*KD cells showed sustained γH2AX foci relative to *ZNF251*WT controls following X-ray exposure, with delayed resolution (Fig. 2A). The cell-cycle inhibitor p21^CIP1/WAF1 was induced at both transcript and protein levels (Fig. 2B–D; Supplementary Fig. 3A, B). In contrast, p16^INK4a was upregulated at the transcript level without a corresponding increase in protein (Fig. 2B and C, D), consistent with the genetic background of MDA-MB-436 cells, which are RB1-null (p.Gly203fs); because p16^INK4a enforces arrest exclusively through the CDK4/6–RB axis, this arm of the senescence program is non-functional in these cells, and senescence-associated growth suppression here depends on p21[31]. p53 protein was also increased (Fig. 2C and D), but because these cells carry a *TP53* frameshift mutation (p.Glu204fs), this likely reflects accumulation of a non-functional product rather than activation of a p53 transcriptional response, indicating that p21 induction in this context is p53-independent[32]. *ZNF251*KD cells also displayed a secretory program: IL-1α, IL-1β, IL-6, IL-8 and TNF-α transcripts were elevated by RT-qPCR, and the corresponding proteins were increased by cytokine array (Fig. 2E and F). To exclude reference-gene artifacts, transcripts were independently normalized to *TBP*, *HPRT1* and *PPIA*, with results concordant with *GAPDH*-normalized measurements (Supplementary Fig. 3C). Two features, however, distinguish this state from classical senescence. First, Lamin B1, whose loss is a defining and largely irreversible feature of terminal senescence[33], was not reduced but rather increased in *ZNF251*KD cells (Fig. 2C, D), indicating that these cells have not entered a stable senescent state. Second, and most importantly, *ZNF251*KD cells failed to undergo stable cell-cycle arrest. Cell-cycle analysis showed only modest G0/G1 accumulation (Fig. 2G) with comparable S-phase fractions (Fig. 2H). Consistent with this, we previously reported that three independent *ZNF251*KD clones grew indistinguishably from *ZNF251*WT cells over 15 days in the absence of drug, in both breast and ovarian cancer backgrounds, and that under olaparib treatment *ZNF251*KD cells maintained a significantly higher Ki-67–positive fraction than *ZNF251*WT cells[9]. Thus, *ZNF251*KD cells continue to cycle even under conditions that suppress proliferation in *ZNF251*-proficient cells. Together, these data define a pseudo-senescent state: cells acquire SA-β-gal positivity, persistent DNA damage, CDK inhibitor induction and a SASP, while retaining Lamin B1 and proliferative capacity[14, 34].

**Figure 2.**
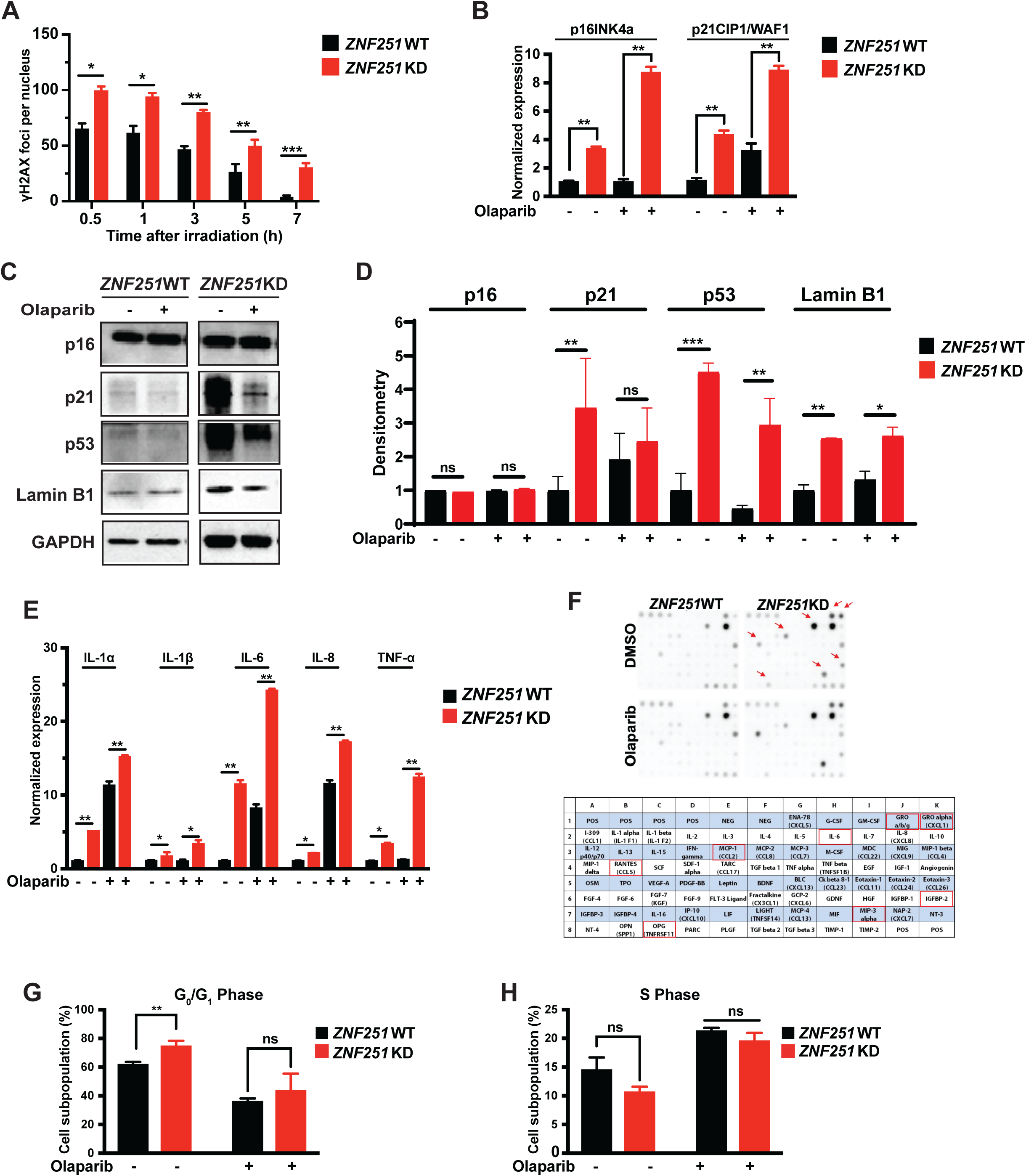
*ZNF251*KD induces pseudo-senescence in *BRCA1*^mut^ breast cancer cells. (A) DNA damage, measured by γH2AX foci formation, was increased in *ZNF251*KD cells compared with *ZNF251*WT cells following X-ray irradiation. (B) Increased expression of canonical senescence regulators p16^INK4a^ and p21^CIP1/WAF1^ in *ZNF251*KD cells, determined by qPCR. (C) Representative Western blots of p16, p21, p53, Lamin B1, and GAPDH (loading control) in *ZNF251*WT and *ZNF251*KD MDA-MB-436 cells treated with vehicle (−) or olaparib (+, 8μM for 3 days). (D) Densitometric quantification of the blots in (C). Band intensities were normalized to GAPDH and expressed relative to vehicle treated *ZNF251*WT cells (set to 1). Black bars, *ZNF251*WT; red bars, *ZNF251*KD. Data are mean ± SD from 3 independent experiments. Statistical significance was determined by two-way ANOVA; horizontal bars indicate significant comparisons (*p<0.05, **p<0.01, ***p<0.001). (E and F) Increased expression of selected senescence-associated secretory phenotype (SASP) factors in *ZNF251*KD breast cancer cells, confirmed by qPCR (E) and cytokine array analysis (F). (G and H) *ZNF251*KD breast cancer cells exhibited increased G0/G1 phase accumulation (G) while maintaining proliferative capacity, as indicated by comparable S-phase populations relative to *ZNF251*WT cells (H). Data represent the means ± SD of three replicates.

### *ZNF251*KD-driven HR hyperactivation promotes DNA replication stress

We previously reported that *ZNF251* deficiency promotes RAD51-dependent HR hyperactivation and PARPi resistance in *BRCA1*^mut^ breast cancer[9]. In the current study, we identify pseudo-senescence as a novel phenotype associated with *ZNF251* loss. We hypothesize that aberrant HR hyperactivation generates replication stress, which contributes to the development of a pseudo-senescent state that enables tumor cells to survive PARPi treatment and acquire resistance. To explore the relationship between HR hyperactivation and replication stress, we first assessed HR repair activity in MDA-MB-436 (*BRCA1*^mut^) cells, including *ZNF251*WT, *ZNF251*KD, and *BRCA1*-reconstituted cells, as *BRCA1* restoration is a well-established mechanism of PARPi resistance in the clinic[4, 35]. Consistent with our previous findings[9], *ZNF251*KD cells exhibited significantly elevated HR activity compared with WT cells, whereas *BRCA1*-reconstituted cells displayed an intermediate level of HR repair activity. Notably, neither *ZNF251*KD nor *BRCA1*-reconstituted cells showed a significant change in HR activity following olaparib treatment. In contrast, olaparib further suppressed HR activity in *ZNF251*WT cells, consistent with their greater sensitivity to PARPi treatment (Fig. 3A). Because HR repair is tightly coordinated with DNA replication, sustained HR hyperactivation may disrupt replication fork homeostasis and induce replication stress, a known trigger of senescence-associated programs[30, 36, 37].

**Figure 3.**
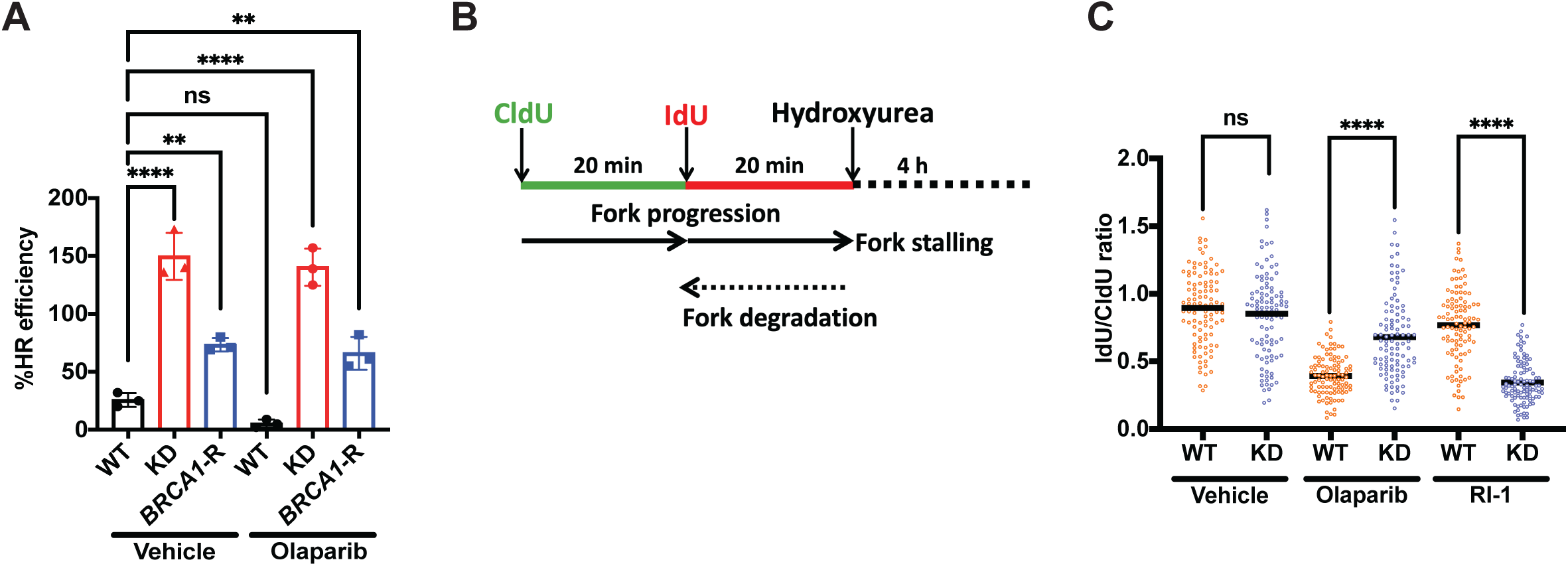
*ZNF251*KD caused replication stress in *BRCA1*^mut^ breast cancer. (A) *ZNF251*KD breast cancer cells exhibit increased HR repair efficiency. (A) Graph summarizing the HR efficiency calculated by the percentage of GFP expression employing the flow cytometry. *ZNF251*WT, *ZNF251*KD, and *BRCA1*-R cells were treated with DMSO or Olaparib. The graph is a representative of 3 independently performed experiments. Significance was calculated with the Mann–Whitney U-test, and ****P < 0.0001 differences between samples. (B) Schematic representation of the protection or degradation of nascent DNA at stalled replication forks employing the DNA fiber assay. (C) Graph summarizing the quantification of the ldU/CIdU ratio for n = 100 DNA fibers analyzed per sample for each experiment (Cells were treated with Olaparib & RI-1). The ratio of IdU/CldU=1 indicates the fork is protected, and the ratio of IdU/CldU<1 indicates the fork is degraded. The graph is representative of 3 independently performed experiments. Significance was calculated with the Mann–Whitney U-test, and the bar indicated is the median for each sample. ****P < 0.0001 differences between samples. Data represent the means ± SD of three replicates.

To determine whether *ZNF251* deficiency alters replication fork dynamics, we performed DNA fiber assays, a single-molecule approach that measures replication fork progression by sequential pulse-labeling of nascent DNA with the thymidine analogs CldU and IdU (Fig. 3B). Under basal conditions, *ZNF251*KD and WT cells exhibited similar IdU/CldU ratios. However, following olaparib treatment, *ZNF251*KD cells displayed a significantly higher IdU/CldU ratio than WT cells, indicating altered replication fork dynamics (Fig. 3C). Importantly, treatment with the RAD51 inhibitor RI-1 attenuated this phenotype, suggesting that RAD51 activity contributes to the replication abnormalities associated with *ZNF251* deficiency. Together with the increased γH2AX foci observed in *ZNF251*KD cells, these findings support a model in which HR hyperactivation promotes replication stress in *ZNF251*-deficient *BRCA1*^mut^ breast cancer cells, potentially contributing to the induction of pseudo-senescence and PARPi resistance.

### Senotherapeutic targeting of (pseudo-)senescence overcomes *ZNF251*KD-mediated PARPi resistance *in vitro*

Cellular senescence can be therapeutically targeted using small-molecule agents collectively termed senotherapeutics, which are broadly categorized into senomorphics and senolytics[13, 38]. Senomorphics modulate the SASP, suppressing its pro-inflammatory effects without eliminating senescent cells, whereas senolytics selectively induce apoptosis in senescent cells by disrupting pro-survival signaling pathways[39, 40]. Given that *ZNF251*KD cells exhibit a senescence-like phenotype, we next asked whether senotherapeutic strategies could overcome PARPi resistance in *BRCA1*-mutant breast cancer cells. We first evaluated senomorphic approaches using inhibitors of pro-inflammatory cytokine signaling, including Bay 11-7082, an NF-κB inhibitor[41], Y-320 (an IL-17A inhibitor)[42], and LMT-28 (an IL-6 receptor antagonist)[43] (Fig. 4A-C). Co-treatment with each of these agents and olaparib significantly reduced cell viability compared with olaparib alone in *ZNF251*KD cells, indicating re-sensitization to olaparib. We then assessed senolytic strategies using BCL-2 family inhibitors. Co-treatment with olaparib and either navitoclax, which targets BCL-2, BCL-XL, and BCL-W, or A-1331852, a selective BCL-XL inhibitor, markedly reduced viability in *ZNF251*KD cells, effectively reversing the olaparib-resistant phenotype (Fig. 4D and E). These findings were recapitulated in *BRCA1*-mutant HCC1937 breast cancer cells (Supplementary Fig. 4). To further validate these findings using a long-term survival assay, we performed clonogenic assays. Consistent with the cell viability results, *ZNF251*KD cells exhibited marked resistance to olaparib compared with *ZNF251*WT cells (Fig. 4F). Importantly, combining olaparib with either BAY 11-7082 or navitoclax markedly reduced clonogenic survival of *ZNF251*KD cells compared with olaparib alone (Fig. 4G and H). Together, these results demonstrate that both senomorphic and senolytic approaches can overcome *ZNF251*KD-mediated PARPi resistance in both short-term cell viability and long-term clonogenic survival assays.

**Figure 4.**
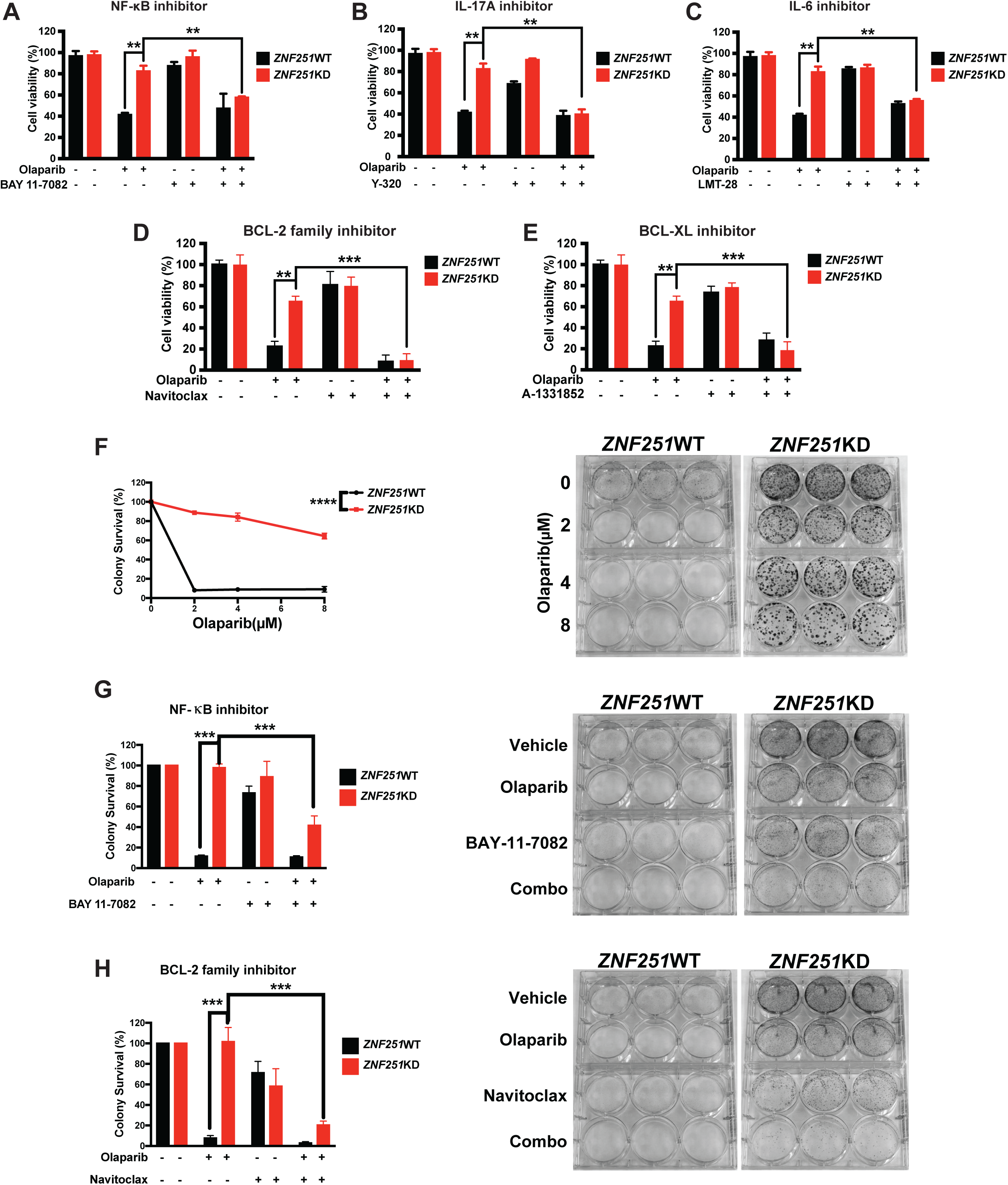
Senotherapeutic targeting of cellular (pseudo-)senescence overcomes *ZNF251*KD-mediated PARPi resistance *in vitro*. (A-C) Cell viability analysis of *ZNF251*WT and *ZNF251*KD breast cancer cells treated with olaparib in combination with the senomorphic agents BAY11-7082 (NF-κB inhibitor) (A), Y-320 (IL-17A inhibitor) (B), or LMT-28 (IL-6 receptor antagonist) (C). Co-treatment with each senomorphic agent significantly reduced the viability of *ZNF251*KD cells compared with olaparib treatment alone, indicating re-sensitization to PARP inhibition. (D and E) Cell viability analysis of *ZNF251*WT and *ZNF251*KD breast cancer cells treated with olaparib in combination with the senolytic agents navitoclax (BCL-2 family inhibitor) (D) or A-1331852 (BCL-XL inhibitor) (E). Co-treatment with each senolytic agent significantly reduced the viability of *ZNF251*KD cells compared with olaparib treatment alone, indicating re-sensitization to PARP inhibition. Data represent the means ± SD of three replicates. (F) Clonogenic survival analysis of *ZNF251*WT and *ZNF251*KD breast cancer cells treated with increasing concentrations of olaparib (0, 2, 4, and 8 μM). Quantification is shown on the left and representative colony images on the right. *ZNF251*KD cells maintained substantially greater clonogenic survival than *ZNF251*WT cells following olaparib treatment, confirming the olaparib-resistant phenotype. (G and H) Clonogenic survival analysis of *ZNF251*WT and *ZNF251*KD cells treated with olaparib alone, BAY 11-7082 (G) or navitoclax (H) alone, or the indicated combinations, with quantification shown on the left and representative colony images on the right. Combination treatment with olaparib and either BAY 11-7082 or navitoclax markedly reduced colony survival of *ZNF251*KD cells compared with olaparib alone.

### Senolytic therapy reverses *ZNF251*KD-induced olaparib resistance *in vivo*

We next assessed whether the BCL-2 family inhibitor navitoclax could reverse *ZNF251*KD-induced PARPi resistance *in vivo*. To test this, 1 × 10 *ZNF251*WT or *ZNF251*KD *BRCA1*^mut^ breast cancer cells were subcutaneously implanted into opposite flanks of 16 female NSG mice. After tumor establishment, approximately 3–4 weeks after implantation, mice were randomized into four treatment groups: vehicle, olaparib, navitoclax, or the combination of olaparib and navitoclax for 28 days (Fig. 5A). Consistent with our in vitro findings, *ZNF251*KD tumors were resistant to olaparib monotherapy, with tumor growth comparable to that of vehicle-treated controls. In contrast, combined treatment with olaparib and navitoclax produced a marked antitumor response, significantly reducing both tumor volume and tumor weight compared with vehicle or either single-agent treatment (Fig. 5B-F). No treatment-related weight loss was observed, supporting the tolerability of both agents (Fig. 5G). These results indicate that BCL-2 family inhibition restores olaparib sensitivity in *ZNF251*-deficient tumors and supports senolytic therapy as a potential strategy to overcome PARPi resistance in *BRCA1*^mut^ breast cancer.

**Figure 5.**
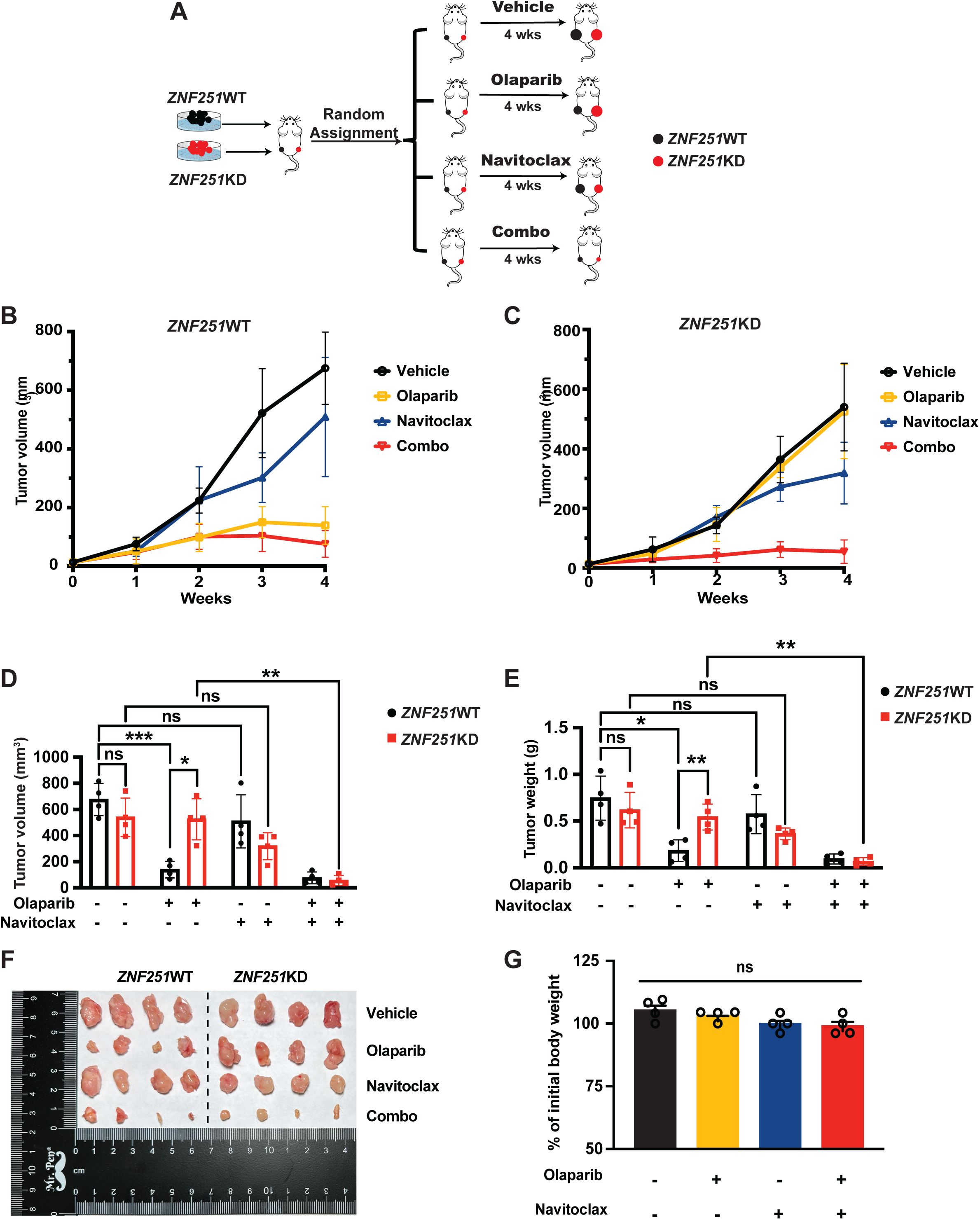
Navitoclax, a BCL-2 family inhibitor, effectively kill *ZNF251*KD olaparib-resistant breast cancer cells in cell-line xenograft (CDX) models. (A) Schematic representation of the CDX *in vivo* study design. (B-E) *In vivo* efficacy analysis in the CDX model showing tumor growth curves (B and C), endpoint tumor volumes (D), and tumor weights (E). Combination treatment with olaparib and navitoclax significantly reduced tumor burden compared with monotherapy treatment groups. (F) Representative images of tumors excised at study endpoint from CDX models treated with vehicle, olaparib, navitoclax, or the combination. Tumors from individual mice are shown; ruler ticks = 1 cm. (G) The combination treatment had no significant effect on recipient body weight. Data represent the means ± SD (n = 4/group).

### BCL-2 family inhibitors improved the sensitivity to olaparib in breast cancer patient samples and effectively restore olaparib sensitivity in olaparib-resistant breast cancer PDX-derived organoids (PDXOs)

To further extend these findings to clinically relevant patient-derived models, we first evaluated the efficacy of BCL-2 family-targeted therapy in primary breast cancer patient-derived cells. Three breast cancer patient-derived cell models with distinct clinical characteristics were treated with increasing concentrations of olaparib in combination with navitoclax. While olaparib monotherapy showed limited efficacy, co-treatment with navitoclax significantly reduced cell viability across all patient-derived models in a dose-dependent manner (Supplementary Fig. 5A-D).

We next evaluated BCL-2 family-targeted therapy in two olaparib-resistant, *BRCA*-mutant triple-negative breast cancer (TNBC) PDX-derived organoids (PDXOs) established from PDX models[25]. PDXO1 carries a germline *BRCA2* frameshift mutation, and PDXO2 carries a somatic *BRCA1* stop-gain mutation (Fig. 6A). To assess homologous recombination (HR) status, we stained for Ki67, γ-H2AX, and RAD51 (Fig. 6B, C). Both PDXOs showed higher Ki67 than control and no detectable RAD51 foci, consistent with HR deficiency, while PDXO1 also showed elevated γ-H2AX. Despite these HR defects, both PDXOs were highly resistant to olaparib monotherapy (Fig. 6D). Navitoclax and A-1331852 alone reduced PDXO viability and further enhanced olaparib sensitivity when combined (data not shown). However, both agents cause dose-limiting thrombocytopenia because BCL-XL is essential for platelet survival[18, 19]. We therefore tested DT2216, a BCL-XL-targeting PROTAC that degrades BCL-XL through VHL, which is poorly expressed in platelets, and thereby spares platelets[21]. DT2216 sensitized both olaparib-resistant PDXOs to olaparib: the DT2216-olaparib combination significantly reduced viability compared with DT2216 alone at every concentration tested (Fig. 6E, F). In PDXO1, DT2216 alone had little effect, yet the combination consistently reduced viability by roughly 30% regardless of dose, indicating that DT2216 sensitizes to olaparib even at the lowest concentration (0.04 µM). In PDXO2, DT2216 alone reduced viability in a dose-dependent manner, and olaparib further enhanced this effect at all doses. Together, these results show that BCL-XL targeting can restore olaparib sensitivity in HR-deficient breast cancer PDXOs.

**Figure 6.**
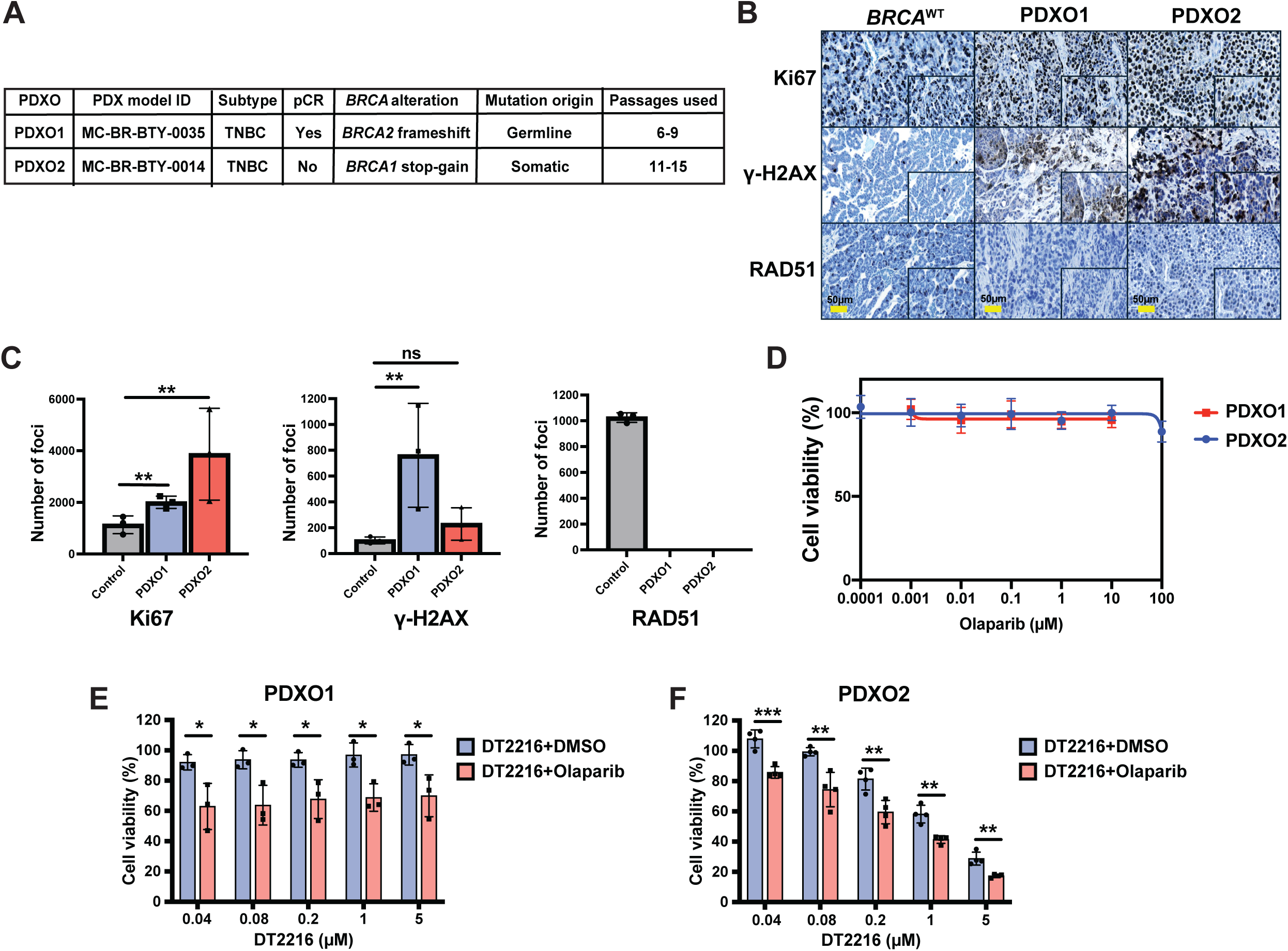
BCL-2 family inhibitor DT2216 restore olaparib sensitivity in olaparib-resistant breast cancer PDX-derived organoids (PDXOs). (A) Characteristics of the two PDXO models. Both are triple-negative breast cancer (TNBC). *BRCA* status was determined by sequencing of the patient’s germline DNA and tumor. PDXO1 (PDX model MC-BR-BTY-0035) carries a germline *BRCA2* frameshift, was derived from a tumor with pCR, and was used at passages 6-9. PDXO2 (MC-BR-BTY-0014) carries a somatic *BRCA1* stop-gain, was derived from a tumor without pCR, and was used at passages 11-15. (B) Representative immunohistochemical staining of Ki67, γ-H2AX, and RAD51 in *BRCA*^WT^ control, PDXO1, and PDXO2. Insets show higher magnification. Scale bars, 50 µm. (C) Quantification of Ki67, γ-H2AX, and RAD51 foci in control, PDXO1, and PDXO2. Both PDXOs show higher Ki67 than control. γ-H2AX is elevated in PDXO1 but not significantly in PDXO2. RAD51 foci are absent in both PDXOs, indicating a homologous recombination defect. Data are mean ± SD. **P < 0.01; ns, not significant. (D) Olaparib dose-response curves for PDXO1 and PDXO2 (0.0001-100 µM). Both organoids are resistant, with viability preserved until the highest dose. (E, F) Cell viability of PDXO1 (E) and PDXO2 (F) treated with increasing concentrations of the BCL-2 family inhibitor DT2216 (0.04-5 µM) combined with DMSO or olaparib 5µM. Adding olaparib significantly reduced viability at every DT2216 dose in both models. Data are mean ± SD, n = 3. *P < 0.05, **P < 0.01, ***P < 0.001.

### *ZNF251*KD redistributes H3K27me3 across the genome

ZNF251 is a KRAB zinc-finger transcription factor, we next first sought to profile its chromatin occupancy. CUT&RUN with the two commercially available ZNF251 antibodies gave no signal enriched over IgG, and public anti-FLAG ChIP-seq of endogenously 3xFLAG-tagged ZNF251 in HepG2 cells (ENCODE) showed a similarly high background. These results suggest that ZNF251 may not bind chromatin stably in the way a typical sequence-specific zinc-finger transcription factor does. Since KRAB domain protein has been implicated in heterochromatin regulation[44], we therefore asked whether loss of ZNF251 changes the chromatin landscape, using CUT&RUN for H3K27me3 and H3K4me3 in ZNF251WT and *ZNF251*KD (clone 3) cells in two independent datasets (RepA and RepB; Fig. 8).

We defined H3K27me3 regions from MACS3 broad peak calls and compared KD with WT after correcting for the global shift in signal. Of these regions, 8,842 gained H3K27me3 beyond the global shift and 7,769 lost it relative to the shift (Fig. 8A). The two classes occupied different chromatin. Regions that gained H3K27me3 were poor in H3K4me3 in both WT and KD cells, whereas regions that lost H3K27me3 were rich in H3K4me3 (Fig. 8A, middle). The ENCODE ZNF251 ChIP-seq track from HepG2 cells was flat across both classes (Fig. 8A, right); this dataset comes from a different cell type and does not exclude ZNF251 binding in MDA-MB-436 cells.

Genome browser views showed the two patterns directly and reproducibly in both datasets. Across a 38 Mb region of chr3, KD cells acquired broad H3K27me3 domains spanning hundreds of kilobases, for example over TAFA1, CNTN3, ROBO1/GBE1, CADM2 and EPHA3, where WT cells had little signal (Fig. 8B). The H3K4me3 and IgG tracks were unchanged across the same region. Across a 25 Mb region of chr12, by contrast, the tall H3K27me3 patches present in WT cells at SLC6A15, CCER1/BTG1, NEDD1 and C12orf42 were flattened in KD cells rather than removed, leaving a lower, more even level of signal (Fig. 8C).

To describe this behavior at the domain scale, we merged blocks of at least 100 kb across gaps of up to 1 Mb to obtain 411 broad H3K27me3 domains. With a threshold of 0.1 CPM, 184 domains gained and 71 lost H3K27me3 in KD cells, and 156 were unchanged (Fig. 8D). Gained domains acquired a broad plateau of H3K27me3 that WT cells lacked (Fig. 8E). Lost domains kept their width, but their tall WT peaks flattened to a lower level rather than disappearing (Fig. 8F). Together, these data show that loss of ZNF251 does not simply raise or lower H3K27me3 but redistributes it: broad new domains form over H3K4me3-poor chromatin while existing focal domains flatten. Senescent cells profoundly remodel their chromatin, most clearly through senescence-associated heterochromatin foci (SAHF)[45], and this global H3K27me3 redistribution is consistent with global remodeling of heterochromatin in ZNF251-depleted cells.

### Other mutations that drive PARPi resistance via HR hyperactivation similarly display features of pseudo-senescence in breast cancer

To determine whether the pseudo-senescence phenotype observed in *ZNF251*KD cells represents a general consequence of HR hyperactivation rather than a gene-or cell line– specific effect, we extended our analysis to additional cancer models with defined defects in HR regulation. Because disruption of the 53BP1-Shieldin pathway is known to restore HR and confer PARPi resistance in *BRCA1*-deficient contexts, we evaluated senescence-associated features in cell lines harboring perturbations in this axis. Specifically, we analyzed *SHLD1*-or *SHLD2*-deficient hTERT RPE-1 cells, a *BRCA1*^WT^, non-transformed retinal pigment epithelial cell line, and MDA-MB-436 cells, a *BRCA1*^mut^ breast cancer cell line. We also examined *53BP1*-deficient U2OS cells, a *BRCA1*^WT^ osteosarcoma cell line.

Across all models, disruption of the 53BP1-Shieldin pathway was consistently associated with increased SA-β-gal activity (Fig. 7A), indicating the presence of a senescence-like phenotype. SHLD1 and SHLD2 are core components of the Shieldin complex, which functions downstream of 53BP1 in non-homologous end joining (NHEJ) to limit DNA end resection; loss of these factors restores HR and promotes PARPi resistance in *BRCA1*-deficient cells[7, 8]. Consistent with this mechanism, *SHLD1*-and *SHLD2*-deficient cells in our study exhibited both olaparib resistance and elevated SA-β-gal activity (Fig. 7B and data not shown). Notably, treatment with senolytic agents, including navitoclax or A-1331852, restored olaparib sensitivity in these models (Fig. 7C-F), supporting a broader role for BCL-2 family inhibition in targeting PARPi resistance associated with pseudo-senescence in breast cancer.

**Figure 7.**
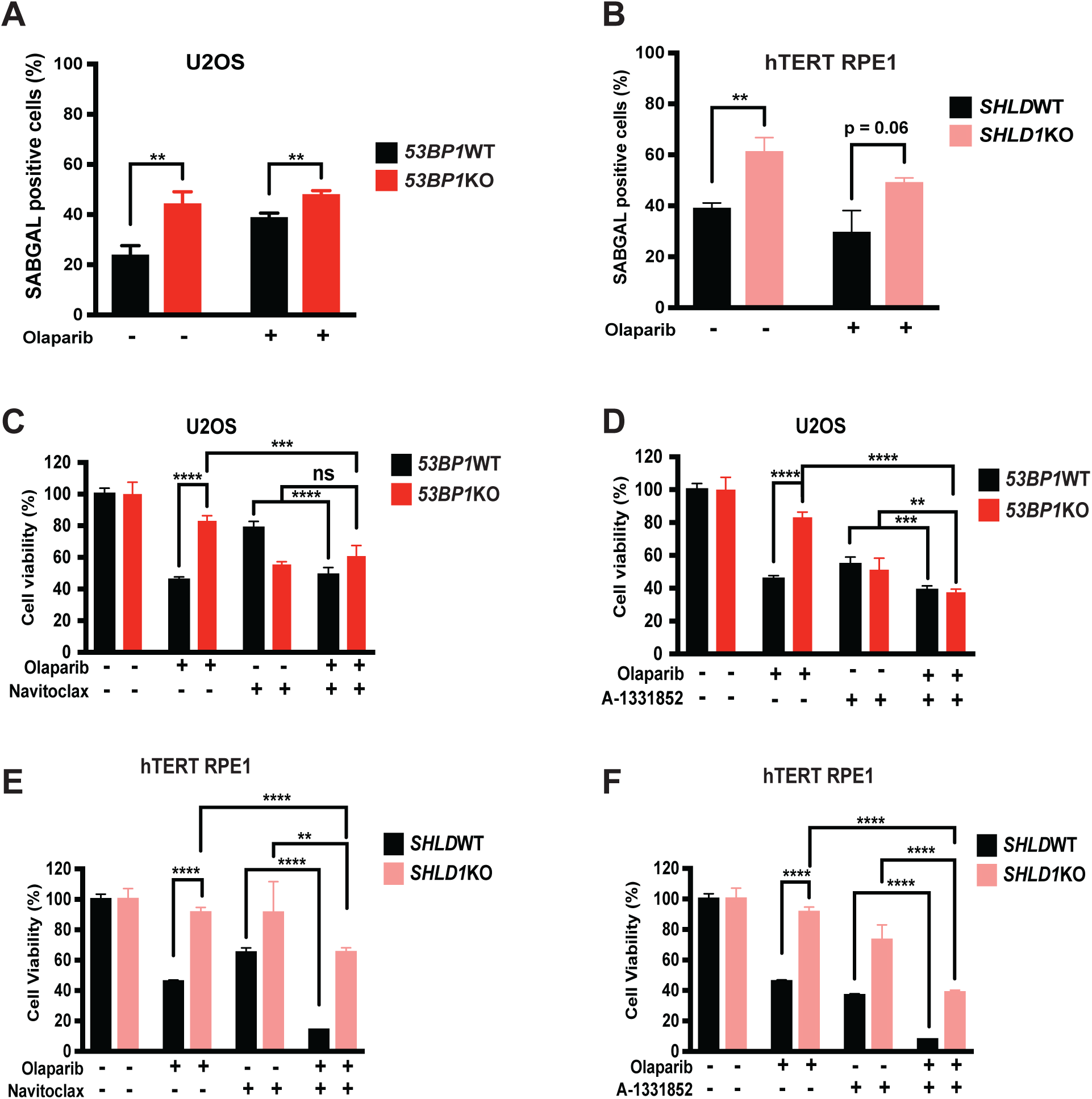
Other mutations that drive PARPi resistance through HR hyperactivation also exhibit features of pseudo-senescence. (A) Quantification of SA-β-gal-positive cells in *53BP1*WT and *53BP1*KO U2OS cells by flow cytometry (%). (B) Quantification of SA-β-gal-positive cells in *SHLD1*WT and *SHLD1*KO hTERT RPE1 cells by flow cytometry (%). (C and D) Cell viability analysis of *53BP1*WT and *53BP1*KO U2OS cells treated with olaparib in combination with the senolytic agents navitoclax (C) or A-1331852 (D). Co-treatment with each senolytic agent significantly reduced the viability of *53BP1*KO cells compared with olaparib treatment alone, indicating re-sensitization to PARP inhibition. (E and F) Cell viability analysis of *SHLD1*WT and *SHLD1*KO hTERT RPE1 cells treated with olaparib in combination with navitoclax (E) or A-1331852 (F). Co-treatment with each senolytic agent significantly reduced the viability of *SHLD1*KO cells compared with olaparib treatment alone, indicating re-sensitization to PARP inhibition. Data represent the means ± SD of three replicates.

Taken together, these data indicate that pseudo-senescence represents a recurrent phenotypic outcome in *BRCA1*-deficient cancer cells that acquire PARPi resistance through HR restoration, rather than a phenotype confined to *ZNF251* deficiency or a single cellular background.

## Discussion

In this study, we identified *ZNF251* deficiency as a driver of RAD51-dependent HR hyperactivation, replication stress, and a pseudo-senescent state that promotes PARPi resistance in *BRCA1*^mut^ breast cancer. PARPis are synthetically lethal in HR-deficient tumors, but resistance often arises through restored HR repair or improved replication fork protection[46, 47]. Our data point to a different route. Loss of *ZNF251* hyperactivates HR, which generates replication stress and gives rise to a pseudo-senescent population that survives PARPi treatment. These cells may be a vulnerability that can be targeted to overcome PARPi resistance (Fig. 9). Our CUT&RUN data show that loss of *ZNF251* is accompanied by a large-scale redistribution of H3K27me3 (Fig. 8), and chromatin compaction is known to influence how DSBs are processed, including end resection and the choice between NHEJ and HR. These observations raise the possibility that ZNF251 restrains HR by regulating chromatin structure, so that its loss relaxes or reorganizes heterochromatin and permits the RAD51-dependent HR hyperactivation we observed.

**Figure 8.**
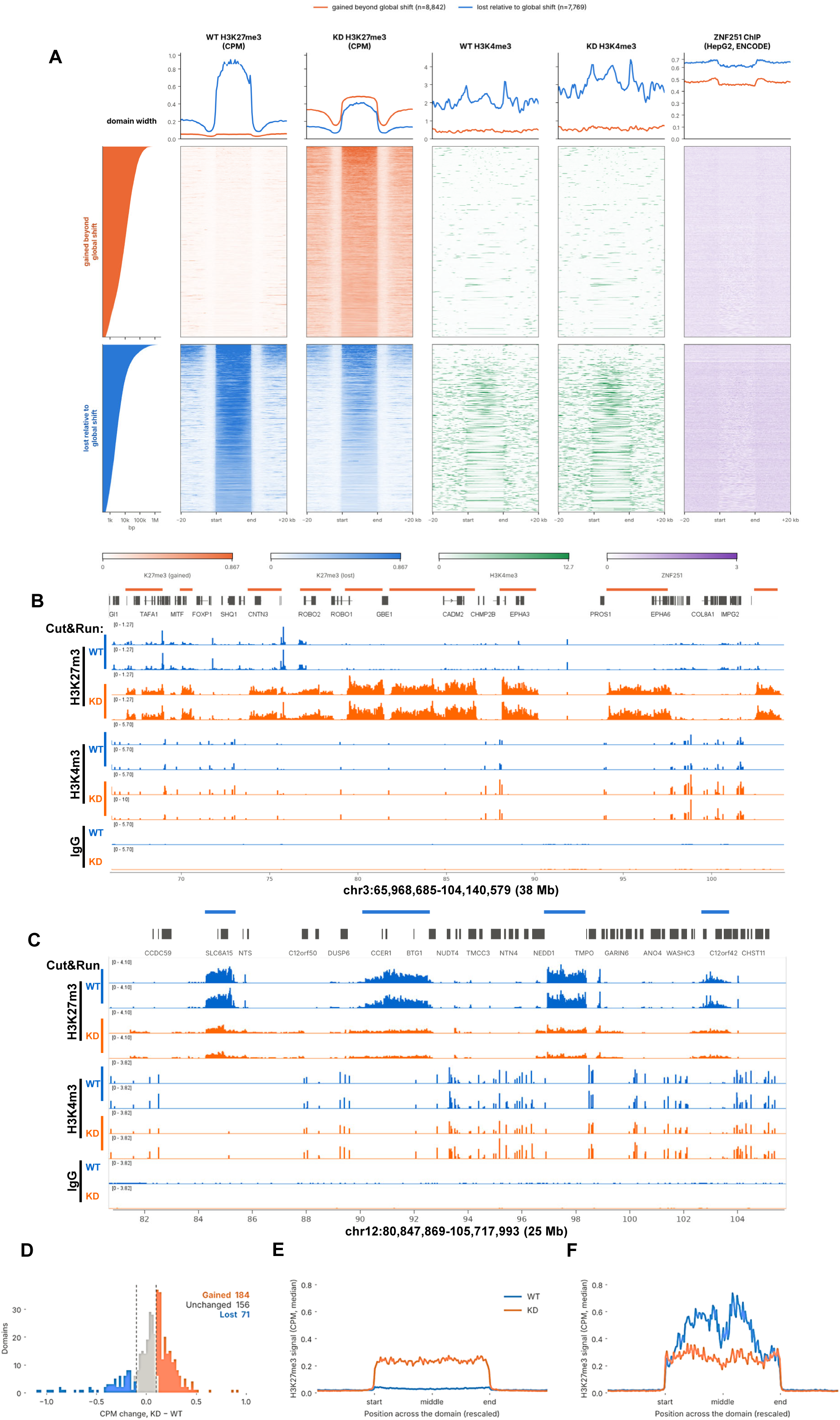
H3K27me3 is redistributed in *ZNF251*-deficient (KD) cells. H3K27me3 regions were defined from MACS3 broad peak calls in two ways. (A) Focal-scale analysis: all called regions, merged within 5 kb, were compared between KD and WT after correcting for the global shift in signal (log2 KD/WT centered on the median). Regions that gained signal beyond the global shift (n = 8,842) or lost signal relative to it (n = 7,769) are shown as heatmaps and mean profiles (WT and KD H3K27me3 and H3K4me3, CPM) with ENCODE HepG2 ZNF251 ChIP-seq for reference (not from MDA-MB-436). Regions are sorted by width; each is scaled to equal length with ±20 kb flanks. (B, C) IGV views of representative gained (chr3) and lost (chr12) regions; bars mark the gained or lost patches. Replicate tracks (RepA, RepB) are shown for WT and KD. (D–F) Domain-scale analysis: blocks of at least 100 kb were merged across gaps of up to 1 Mb, which ignores small interruptions, to give 411 broad domains for studying the overall gain or loss trend. (D) Change in mean H3K27me3 signal (CPM, KD − WT) per domain; domains are classed as gained (> +0.1 CPM, n = 184), unchanged (n = 156) or lost (< −0.1 CPM, n = 71). (E, F) Median H3K27me3 profile across gained (E) and lost (F) domains, each rescaled to equal length (WT blue, KD orange); lost domains flatten but do not widen. The two region sets answer different questions, and gain and loss are defined relative to the global shift, not as absolute counts. KD is clone 3.

**Figure 9.**
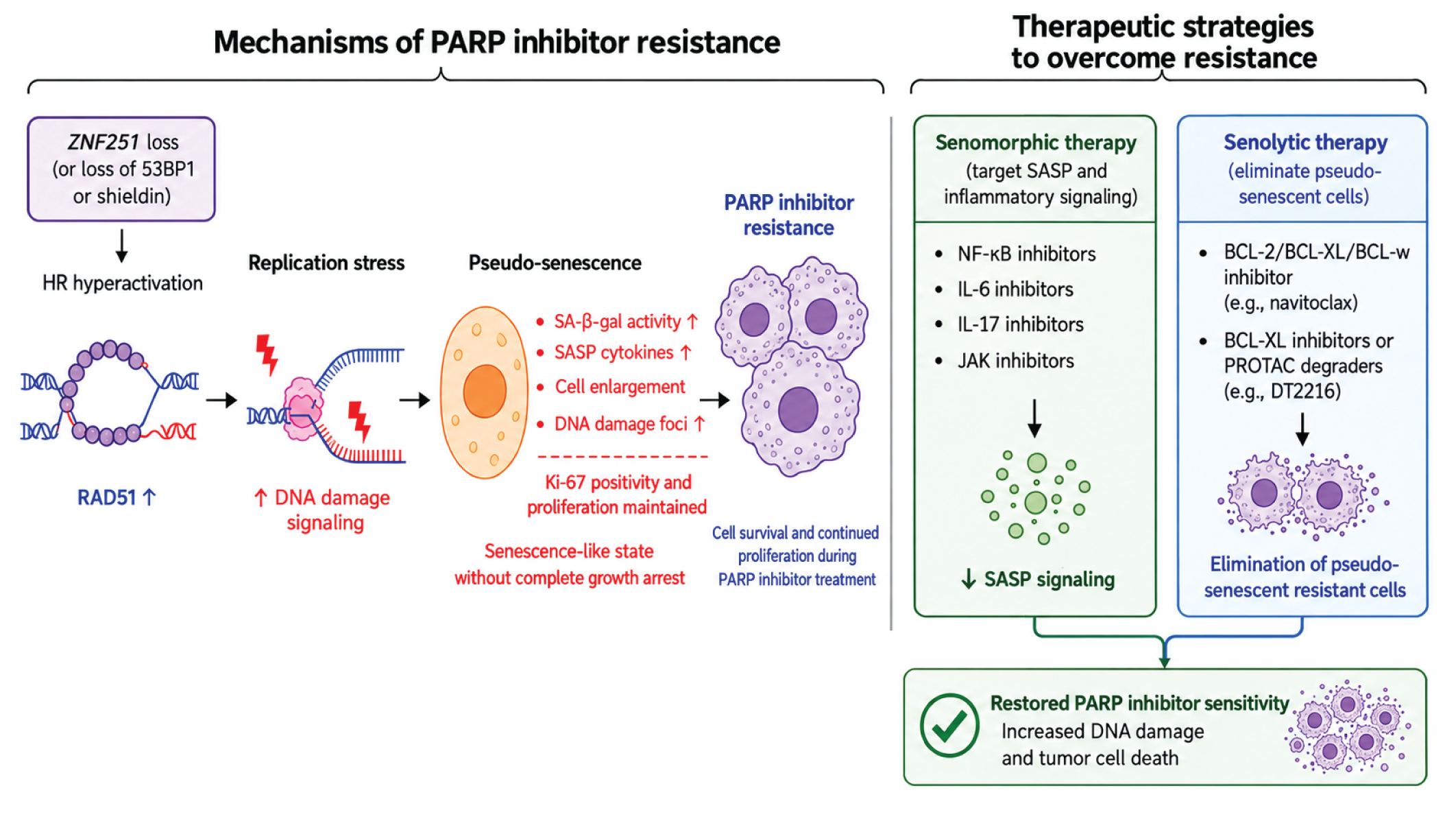
Schematic illustration of the mechanisms by which *ZNF251* deficiency and *53BP1* or *Shieldin* loss drive PARPi resistance in breast cancer and the therapeutic strategies to overcome this resistance. *ZNF251* deficiency, or loss of *53BP1*/*Shieldin*, promotes HR hyperactivation, replication stress, and pseudo-senescence, allowing cells to survive PARPi treatment and continue proliferating. Senomorphic or senolytic therapies can target these resistant cells, restore PARPi sensitivity, and increase tumor cell death.

The main new finding is that *ZNF251* deficiency induces pseudo-senescence, an incomplete and reversible senescence-like state. *ZNF251*-deficient cells showed canonical senescence features, including higher SA-β-gal activity and more γH2AX foci, but they did not arrest durably and kept their capacity to proliferate. The expanded marker panel shows which parts of the senescence program are active. p21^CIP1/WAF1 was induced at both the transcript and protein levels, whereas p16^INK4a was up only at the transcript level. MDA-MB-436 cells are RB1-null (p.Gly203fs), so the p16–RB arm cannot be executed in this background and growth suppression relies on p21 alone. Lamin B1, whose loss is a defining and largely irreversible feature of terminal senescence, was increased rather than reduced in *ZNF251*KD cells. Retained Lamin B1, incomplete arrest, and a nonfunctional p16–RB axis set this state apart from classical senescence and may explain why the cells never settle into stable arrest. We propose that replication stress–induced pseudo-senescence lets cancer cells adapt to chronic DNA damage, avoid PARPi cytotoxicity, and resume proliferation after treatment. The state was not specific to *ZNF251* loss. We also saw it in other PARPi-resistant models, including *53BP1*-deficient and *Shieldin*-deficient cells, so senescence-like plasticity may be a broader and underrecognized route to PARPi resistance. As with therapy-induced senescence, these cells may persist through SASP-associated inflammatory and pro-survival signaling and by escaping stable growth arrest[15, 34, 48].

Pseudo-senescence also creates a therapeutic vulnerability[39, 40]. Senotherapeutics overcame *ZNF251*KD-mediated PARPi resistance, which suggests these cells rely on senescence-associated survival pathways. The BCL-2/BCL-XL inhibitor navitoclax and the selective BCL-XL inhibitor A-1331852 reversed olaparib resistance in vivo and potently eliminated breast cancer patient-derived cells. This sensitivity probably reflects a greater dependence on anti-apoptotic proteins such as BCL-XL and MCL-1, possibly reinforced by persistent replication stress and DNA damage response signaling[49, 50]. The results support pairing PARPis with senolytics to kill resistant cells and keep a pseudo-senescent reservoir from persisting.

Whether ZNF251 itself can be targeted is a fair question, and direct targeting looks difficult. ZNF251 is a KRAB-domain zinc finger transcription factor, a class that lacks enzymatic activity and well-defined ligand-binding pockets. Resistance here also comes from loss of function, so inhibiting ZNF251 would be expected to worsen the phenotype rather than reverse it. We favor an indirect strategy that targets the downstream state created by *ZNF251* loss. This fits the broader principle that resistance adaptations create new dependencies[51]. *BRCA1*-mutant tumors that become PARPi-resistant through 53BP1 or Shieldin loss are sensitive to ATR or Polθ inhibition, and here, cells in a pseudo-senescent state depend on BCL-2 family proteins. Other efforts to overcome PARPi resistance have targeted HR regulators rather than the resistance lesion, including NAT10[52] and Jab1/COPS5 in triple-negative breast cancer[53]. In this framework, *ZNF251* status would act as a predictive biomarker for senolytic combinations rather than as a drug target, and clinical-stage agents such as navitoclax and the BCL-XL– directed PROTAC DT2216[21] make this feasible. Translation faces three main challenges: on-target thrombocytopenia from BCL-XL inhibition, the lack of assays that reliably identify pseudo-senescent tumor cells in patient material, and the optimal scheduling of PARPi and senolytic agents.

Our study has limitations. Our *in vivo* experiments used subcutaneous flank implantation instead of orthotopic mammary fat pad inoculation. Orthotopic models better reflect the mammary microenvironment and its effects on drug penetrance, tissue architecture, and senescence dynamics. We chose the flank for reproducible tumor measurement and consistency with our prior study, but without the native stroma, both the size of the senolytic effect and the makeup of the SASP could differ. NSG mice also lack functional T, B, and NK cells. Senescent cells are normally cleared through SASP-mediated immune surveillance, which our model cannot capture, so the senolytic efficacy we observed reflects direct, cell-autonomous killing rather than immune clearance. In an immunocompetent host, SASP-driven immune recruitment could either improve clearance of pseudo-senescent cells or fuel tumor-supportive inflammation, and immunocompetent models are needed to tell which. MDA-MB-436 cells came from a pleural effusion rather than a primary tumor. They have therefore been selected *in vivo* for metastatic fitness and may not fully reflect stress responses in earlier-stage disease. We partly addressed this by confirming key findings in HCC1937 cells and in PDX-derived organoids, but models derived from primary *BRCA1*^mut^ lesions would strengthen generalizability. Finally, we studied only *BRCA1*^mut^ and *BRCA1*-methylated contexts. Because the proposed mechanism depends on HR hyperactivation and replication stress rather than on *BRCA1* itself, it may extend to HR deficiency more broadly, including *BRCA2*^mut^ breast cancer and high-grade serous ovarian carcinoma, where PARPis are standard of care. Whether *ZNF251*-associated resistance strictly requires *BRCA1* deficiency or is general to HR-deficient tumors remains open.

In summary, *ZNF251* loss promotes RAD51-dependent HR hyperactivation, which causes replication stress and a pseudo-senescent state that lets *BRCA1*^mut^ breast cancer cells survive PARPi treatment. Targeting this state with senotherapeutics, particularly BCL-2 family inhibitors, may eliminate resistant cells and make PARPi responses last longer in the clinic. Next steps are to define how *ZNF251* loss drives HR hyperactivation, replication stress, and pseudo-senescence; test the long-term efficacy and safety of PARPi–senolytic combinations in orthotopic and immunocompetent models; extend the work to *BRCA2*^mut^ and ovarian cancer models; and develop biomarkers for tumors enriched in pseudo-senescent cells. Future studies should also assess *ZNF251* expression in patient tumor tissue and test its association with PARPi response and clinical outcomes in larger, well-annotated cohorts.

## Supporting information

Combined Supplementary Figures

## Authors’ Contributions

Conception and design: J. Huang

**Development of methodology:** Z. Tian, S. Chatla, S. Indulkar, L. Chen, J. Huang

**Acquisition of data (provided animals, acquired and managed patients, provided facilities, etc.):** Z. Tian, S. Chatla, S. Indulkar, Y. Liu, A. Morris, B. V Le, D. Kim, XY. Wei, Y. L, D. Yang. L. Chen

**Analysis and interpretation of data (e.g., statistical analysis, biostatistics, computational analysis):** Z. Tian, S. Chatla, A. Pompetti, G. Calendo, L. Chen

**Writing, review, and/or revision of the manuscript:** S. Chatla, Z. Tian, L. Chen, J. Huang

**Administrative, technical, or material support (i.e., reporting or organizing Data, constructing databases):** C. Wang, T. Edmonston, Z. Lou, L. Chen, T. Skorski, L. Wang, J. Huang

**Study supervision:** L. Chen, T. Skorski, L. Wang, J. Huang

## Acknowledgments

We sincerely thank Dr. Peter S. Klein at the University of Pennsylvania; Drs. Jean-Pierre Issa, Xiaoxin (Luke) Chen, Nora Engel, Shumei Song, and Jaroslav Jelinek at the Coriell Institute for Medical Research; and Dr. Chris Lord at The Institute of Cancer Research, London, for their insightful comments and discussions. We sincerely thank Diego A. Morales Scheihing for his assistance with the experiments involving patient samples. We also thank all members of the Huang, Wang, Skorski and Chen laboratories for their help and valuable discussions. We specifically thank Dr. Xue Wang at the Mayo Clinic for her support with immunohistochemistry (IHC) staining of patient-derived organoids (PDXOs).

## References

[1] C. Underhill, M. Toulmonde, H. Bonnefoi, A review of PARP inhibitors: from bench to bedside, Ann Oncol, 22 (2011) 268–279.

[2] L. Cortesi, H.S. Rugo, C. Jackisch, An Overview of PARP Inhibitors for the Treatment of Breast Cancer, Target Oncol, 16 (2021) 255–282.

[3] M. Robson, S.A. Im, E. Senkus, B. Xu, S.M. Domchek, N. Masuda, S. Delaloge, W. Li, N. Tung, A. Armstrong, W. Wu, C. Goessl, S. Runswick, P. Conte, Olaparib for Metastatic Breast Cancer in Patients with a Germline BRCA Mutation, N Engl J Med, 377 (2017) 523–533.

[4] H. Li, Z.Y. Liu, N. Wu, Y.C. Chen, Q. Cheng, J. Wang, PARP inhibitor resistance: the underlying mechanisms and clinical implications, Mol Cancer, 19 (2020) 107.

[5] S.M. Noordermeer, H. van Attikum, PARP Inhibitor Resistance: A Tug-of-War in BRCA-Mutated Cells, Trends Cell Biol, 29 (2019) 820–834.

[6] P.C. Fong, T.A. Yap, D.S. Boss, C.P. Carden, M. Mergui-Roelvink, C. Gourley, J. De Greve, J. Lubinski, S. Shanley, C. Messiou, R. A’Hern, A. Tutt, A. Ashworth, J. Stone, J. Carmichael, J.H. Schellens, J.S. de Bono, S.B. Kaye, Poly(ADP)-ribose polymerase inhibition: frequent durable responses in BRCA carrier ovarian cancer correlating with platinum-free interval, J Clin Oncol, 28 (2010) 2512–2519.

[7] J.E. Jaspers, A. Kersbergen, U. Boon, W. Sol, L. van Deemter, S.A. Zander, R. Drost, E. Wientjens, J. Ji, A. Aly, J.H. Doroshow, A. Cranston, N.M. Martin, A. Lau, M.J. O’Connor, S. Ganesan, P. Borst, J. Jonkers, S. Rottenberg, Loss of 53BP1 causes PARP inhibitor resistance in Brca1-mutated mouse mammary tumors, Cancer Discov, 3 (2013) 68–81.

[8] H. Dev, T.W. Chiang, C. Lescale, I. de Krijger, A.G. Martin, D. Pilger, J. Coates, M. Sczaniecka-Clift, W. Wei, M. Ostermaier, M. Herzog, J. Lam, A. Shea, M. Demir, Q. Wu, F. Yang, B. Fu, Z. Lai, G. Balmus, R. Belotserkovskaya, V. Serra, M.J. O’Connor, A. Bruna, P. Beli, L. Pellegrini, C. Caldas, L. Deriano, J.J.L. Jacobs, Y. Galanty, S.P. Jackson, Shieldin complex promotes DNA end-joining and counters homologous recombination in BRCA1-null cells, Nat Cell Biol, 20 (2018) 954–965.

[9] H. Li, S. Chatla, X. Liu, Z. Tian, U. Vekariya, P. Wang, D. Kim, S. Octaviani, Z. Lian, G. Morton, Z. Feng, D. Yang, K. Sullivan-Reed, W. Childers, X. Yu, K.N. Chitrala, J. Madzo, T. Skorski, J. Huang, ZNF251 haploinsufficiency confers PARP inhibitors resistance in BRCA1-mutated cancer cells through activation of homologous recombination, Cancer Lett, 613 (2025) 217505.

[10] V. Gorgoulis, P.D. Adams, A. Alimonti, D.C. Bennett, O. Bischof, C. Bishop, J. Campisi, M. Collado, K. Evangelou, G. Ferbeyre, J. Gil, E. Hara, V. Krizhanovsky, D. Jurk, A.B. Maier, M. Narita, L. Niedernhofer, J.F. Passos, P.D. Robbins, C.A. Schmitt, J. Sedivy, K. Vougas, T. von Zglinicki, D. Zhou, M. Serrano, M. Demaria, Cellular Senescence: Defining a Path Forward, Cell, 179 (2019) 813–827.

[11] R. Kumari, P. Jat, Mechanisms of Cellular Senescence: Cell Cycle Arrest and Senescence Associated Secretory Phenotype, Front Cell Dev Biol, 9 (2021) 645593.

[12] B. Wang, J. Han, J.H. Elisseeff, M. Demaria, The senescence-associated secretory phenotype and its physiological and pathological implications, Nat Rev Mol Cell Biol, 25 (2024) 958–978.

[13] L. Zhang, L.E. Pitcher, V. Prahalad, L.J. Niedernhofer, P.D. Robbins, Targeting cellular senescence with senotherapeutics: senolytics and senomorphics, FEBS J, 290 (2023) 1362–1383.

[14] I. Sreeram, S. Plans-Marin, M. Cruz-Rodriguez, E. Aliagas, D. Palau-Gallinat, C. Munoz-Pinedo, E. Nadal, Pseudo-senescence induced by palbociclib does not sensitise pleural mesothelioma cells to combinations with senolytics, Cell Death Dis, 17 (2026).

[15] C.A. Schmitt, B. Wang, M. Demaria, Senescence and cancer -role and therapeutic opportunities, Nat Rev Clin Oncol, 19 (2022) 619–636.

[16] M. Paez-Ribes, E. Gonzalez-Gualda, G.J. Doherty, D. Munoz-Espin, Targeting senescent cells in translational medicine, EMBO Mol Med, 11 (2019) e10234.

[17] N.N. Mohamad Anuar, N.S. Nor Hisam, S.L. Liew, A. Ugusman, Clinical Review: Navitoclax as a Pro-Apoptotic and Anti-Fibrotic Agent, Front Pharmacol, 11 (2020) 564108.

[18] L. Wang, G.A. Doherty, A.S. Judd, Z.F. Tao, T.M. Hansen, R.R. Frey, X. Song, M. Bruncko, A.R. Kunzer, X. Wang, M.D. Wendt, J.A. Flygare, N.D. Catron, R.A. Judge, C.H. Park, S. Shekhar, D.C. Phillips, P. Nimmer, M.L. Smith, S.K. Tahir, Y. Xiao, J. Xue, H. Zhang, P.N. Le, M.J. Mitten, E.R. Boghaert, W. Gao, P. Kovar, E.F. Choo, D. Diaz, W.J. Fairbrother, S.W. Elmore, D. Sampath, J.D. Leverson, A.J. Souers, Discovery of A-1331852, a First-in-Class, Potent, and Orally-Bioavailable BCL-X(L) Inhibitor, ACS Med Chem Lett, 11 (2020) 1829–1836.

[19] W.H. Wilson, O.A. O’Connor, M.S. Czuczman, A.S. LaCasce, J.F. Gerecitano, J.P. Leonard, A. Tulpule, K. Dunleavy, H. Xiong, Y.L. Chiu, Y. Cui, T. Busman, S.W. Elmore, S.H. Rosenberg, A.P. Krivoshik, S.H. Enschede, R.A. Humerickhouse, Navitoclax, a targeted high-affinity inhibitor of BCL-2, in lymphoid malignancies: a phase 1 dose-escalation study of safety, pharmacokinetics, pharmacodynamics, and antitumour activity, Lancet Oncol, 11 (2010) 1149–1159.

[20] T. Oltersdorf, S.W. Elmore, A.R. Shoemaker, R.C. Armstrong, D.J. Augeri, B.A. Belli, M. Bruncko, T.L. Deckwerth, J. Dinges, P.J. Hajduk, M.K. Joseph, S. Kitada, S.J. Korsmeyer, A.R. Kunzer, A. Letai, C. Li, M.J. Mitten, D.G. Nettesheim, S. Ng, P.M. Nimmer, J.M. O’Connor, A. Oleksijew, A.M. Petros, J.C. Reed, W. Shen, S.K. Tahir, C.B. Thompson, K.J. Tomaselli, B. Wang, M.D. Wendt, H. Zhang, S.W. Fesik, S.H. Rosenberg, An inhibitor of Bcl-2 family proteins induces regression of solid tumours, Nature, 435 (2005) 677–681.

[21] Y. He, R. Koch, V. Budamagunta, P. Zhang, X. Zhang, S. Khan, D. Thummuri, Y.T. Ortiz, X. Zhang, D. Lv, J.S. Wiegand, W. Li, A.C. Palmer, G. Zheng, D.M. Weinstock, D. Zhou, DT2216-a Bcl-xL-specific degrader is highly active against Bcl-xL-dependent T cell lymphomas, J Hematol Oncol, 13 (2020) 95.

[22] T. Saleh, V.J. Carpenter, L. Tyutyunyk-Massey, G. Murray, J.D. Leverson, A.J. Souers, M.R. Alotaibi, A.C. Faber, J. Reed, H. Harada, D.A. Gewirtz, Clearance of therapy-induced senescent tumor cells by the senolytic ABT-263 via interference with BCL-X(L) -BAX interaction, Mol Oncol, 14 (2020) 2504–2519.

[23] A. Shahbandi, S.G. Rao, A.Y. Anderson, W.D. Frey, J.O. Olayiwola, N.A. Ungerleider, J.G. Jackson, BH3 mimetics selectively eliminate chemotherapy-induced senescent cells and improve response in TP53 wild-type breast cancer, Cell Death Differ, 27 (2020) 3097–3116.

[24] A. Softah, M.R. Alotaibi, A.R. Alhoshani, T. Saleh, K. Alhazzani, M.M. Almutairi, R. AlRowis, S. Alshehri, N.A. Albekairy, H. Harada, R. Boyd, E. Chakraborty, D.A. Gewirtz, H.M. As Sobeai, The Combination of Radiation with PARP Inhibition Enhances Senescence and Sensitivity to the Senolytic, Navitoclax, in Triple Negative Breast Tumor Cells, Biomedicines, 11 (2023).

[25] M.P. Goetz, K.R. Kalari, V.J. Suman, A.M. Moyer, J. Yu, D.W. Visscher, T.J. Dockter, P.T. Vedell, J.P. Sinnwell, X. Tang, K.J. Thompson, S.A. McLaughlin, A. Moreno-Aspitia, J.A. Copland, D.W. Northfelt, R.J. Gray, K. Hunt, A. Conners, H. Sicotte, J.E. Eckel-Passow, J.P. Kocher, J.N. Ingle, M.S. Ellingson, M. McDonough, E.D. Wieben, R. Weinshilboum, L. Wang, J.C. Boughey, Tumor Sequencing and Patient-Derived Xenografts in the Neoadjuvant Treatment of Breast Cancer, J Natl Cancer Inst, 109 (2017).

[26] U. Vekariya, M. Toma, M. Nieborowska-Skorska, B.V. Le, M.C. Caron, A.M. Kukuyan, K. Sullivan-Reed, P. Podszywalow-Bartnicka, K.N. Chitrala, J. Atkins, M. Drzewiecka, W. Feng, J. Chan, S. Chatla, K. Golovine, J. Jelinek, T. Sliwinski, J. Ghosh, K. Matlawska-Wasowska, G. Chandramouly, R. Nejati, M. Wasik, S.M. Sykes, K. Piwocka, E. Hadzijusufovic, P. Valent, R.T. Pomerantz, G. Morton, W. Childers, H. Zhao, E.M. Paietta, R.L. Levine, M.S. Tallman, H.F. Fernandez, M.R. Litzow, G.P. Gupta, J.Y. Masson, T. Skorski, DNA polymerase theta protects leukemia cells from metabolically induced DNA damage, Blood, 141 (2023) 2372–2389.

[27] S. Maifrede, B.V. Le, M. Nieborowska-Skorska, K. Golovine, K. Sullivan-Reed, W.M.B. Dunuwille, J. Nacson, M. Hulse, K. Keith, J. Madzo, L.B. Caruso, Z. Gazze, Z. Lian, A. Padella, K.N. Chitrala, B.A. Bartholdy, K. Matlawska-Wasowska, D. Di Marcantonio, G. Simonetti, G. Greiner, S.M. Sykes, P. Valent, E.M. Paietta, M.S. Tallman, H.F. Fernandez, M.R. Litzow, M.D. Minden, J. Huang, G. Martinelli, G.S. Vassiliou, I. Tempera, K. Piwocka, N. Johnson, G.A. Challen, T. Skorski, TET2 and DNMT3A Mutations Exert Divergent Effects on DNA Repair and Sensitivity of Leukemia Cells to PARP Inhibitors, Cancer Res, 81 (2021) 5089–5101.

[28] A.P. Lombard, C.M. Armstrong, L.S. D’Abronzo, S. Ning, A.R. Leslie, M. Sharifi, W. Lou, C.P. Evans, M. Dall’Era, H.W. Chen, X. Chen, A.C. Gao, Olaparib-Induced Senescence Is Bypassed through G2-M Checkpoint Override in Olaparib-Resistant Prostate Cancer, Mol Cancer Ther, 21 (2022) 677–685.

[29] X. Zhang, J. Yao, X. Li, N. Niu, Y. Liu, R.A. Hajek, G. Peng, S. Westin, A.K. Sood, J. Liu, Targeting polyploid giant cancer cells potentiates a therapeutic response and overcomes resistance to PARP inhibitors in ovarian cancer, Sci Adv, 9 (2023) eadf7195.

[30] F. Rodier, J.P. Coppe, C.K. Patil, W.A. Hoeijmakers, D.P. Munoz, S.R. Raza, A. Freund, E. Campeau, A.R. Davalos, J. Campisi, Persistent DNA damage signalling triggers senescence-associated inflammatory cytokine secretion, Nat Cell Biol, 11 (2009) 973–979.

[31] A. Takahashi, N. Ohtani, E. Hara, Irreversibility of cellular senescence: dual roles of p16INK4a/Rb-pathway in cell cycle control, Cell Div, 2 (2007) 10.

[32] T. Matsuda, T. Kato, K. Kiyotani, Y.E. Tarhan, V. Saloura, S. Chung, K. Ueda, Y. Nakamura, J.H. Park, p53-independent p21 induction by MELK inhibition, Oncotarget, 8 (2017) 57938–57947.

[33] A. Freund, R.M. Laberge, M. Demaria, J. Campisi, Lamin B1 loss is a senescence-associated biomarker, Mol Biol Cell, 23 (2012) 2066–2075.

[34] T. Saleh, L. Tyutyunyk-Massey, D.A. Gewirtz, Tumor Cell Escape from Therapy-Induced Senescence as a Model of Disease Recurrence after Dormancy, Cancer Res, 79 (2019) 1044–1046.

[35] L.M. Jackson, G.L. Moldovan, Mechanisms of PARP1 inhibitor resistance and their implications for cancer treatment, NAR Cancer, 4 (2022) zcac042.

[36] R. Di Micco, M. Fumagalli, A. Cicalese, S. Piccinin, P. Gasparini, C. Luise, C. Schurra, M. Garre, P.G. Nuciforo, A. Bensimon, R. Maestro, P.G. Pelicci, F. d’Adda di Fagagna, Oncogene-induced senescence is a DNA damage response triggered by DNA hyper-replication, Nature, 444 (2006) 638–642.

[37] J. Bartkova, N. Rezaei, M. Liontos, P. Karakaidos, D. Kletsas, N. Issaeva, L.V. Vassiliou, E. Kolettas, K. Niforou, V.C. Zoumpourlis, M. Takaoka, H. Nakagawa, F. Tort, K. Fugger, F. Johansson, M. Sehested, C.L. Andersen, L. Dyrskjot, T. Orntoft, J. Lukas, C. Kittas, T. Helleday, T.D. Halazonetis, J. Bartek, V.G. Gorgoulis, Oncogene-induced senescence is part of the tumorigenesis barrier imposed by DNA damage checkpoints, Nature, 444 (2006) 633–637.

[38] T. Saliev, P.B. Singh, Targeting Senescence: A Review of Senolytics and Senomorphics in Anti-Aging Interventions, Biomolecules, 15 (2025).

[39] J.L. Kirkland, T. Tchkonia, Cellular Senescence: A Translational Perspective, EBioMedicine, 21 (2017) 21–28.

[40] L.J. Niedernhofer, P.D. Robbins, Senotherapeutics for healthy ageing, Nat Rev Drug Discov, 17 (2018) 377.

[41] J. Lee, M.H. Rhee, E. Kim, J.Y. Cho, BAY 11-7082 is a broad-spectrum inhibitor with anti-inflammatory activity against multiple targets, Mediators Inflamm, 2012 (2012) 416036.

[42] H. Ushio, S. Ishibuchi, K. Oshita, N. Seki, H. Kataoka, K. Sugahara, K. Adachi, K. Chiba, A new phenylpyrazoleanilide, y-320, inhibits interleukin 17 production and ameliorates collagen-induced arthritis in mice and cynomolgus monkeys, Pharmaceuticals (Basel), 7 (2013) 1–17.

[43] S.S. Hong, J.H. Choi, S.Y. Lee, Y.H. Park, K.Y. Park, J.Y. Lee, J. Kim, V. Gajulapati, J.I. Goo, S. Singh, K. Lee, Y.K. Kim, S.H. Im, S.H. Ahn, S. Rose-John, T.H. Heo, Y. Choi, A Novel Small-Molecule Inhibitor Targeting the IL-6 Receptor beta Subunit, Glycoprotein 130, J Immunol, 195 (2015) 237–245.

[44] D.C. Schultz, K. Ayyanathan, D. Negorev, G.G. Maul, F.J. Rauscher, 3rd, SETDB1: a novel KAP-1-associated histone H3, lysine 9-specific methyltransferase that contributes to HP1-mediated silencing of euchromatic genes by KRAB zinc-finger proteins, Genes Dev, 16 (2002) 919–932.

[45] M. Narita, S. Nunez, E. Heard, M. Narita, A.W. Lin, S.A. Hearn, D.L. Spector, G.J. Hannon, S.W. Lowe, Rb-mediated heterochromatin formation and silencing of E2F target genes during cellular senescence, Cell, 113 (2003) 703–716.

[46] C.J. Lord, A. Ashworth, PARP inhibitors: Synthetic lethality in the clinic, Science, 355 (2017) 1152–1158.

[47] O. Kondrashova, M. Nguyen, K. Shield-Artin, A.V. Tinker, N.N.H. Teng, M.I. Harrell, M.J. Kuiper, G.Y. Ho, H. Barker, M. Jasin, R. Prakash, E.M. Kass, M.R. Sullivan, G.J. Brunette, K.A. Bernstein, R.L. Coleman, A. Floquet, M. Friedlander, G. Kichenadasse, D.M. O’Malley, A. Oza, J. Sun, L. Robillard, L. Maloney, D. Bowtell, H. Giordano, M.J. Wakefield, S.H. Kaufmann, A.D. Simmons, T.C. Harding, M. Raponi, I.A. McNeish, E.M. Swisher, K.K. Lin, C.L. Scott, A.S. Group, Secondary Somatic Mutations Restoring RAD51C and RAD51D Associated with Acquired Resistance to the PARP Inhibitor Rucaparib in High-Grade Ovarian Carcinoma, Cancer Discov, 7 (2017) 984–998.

[48] S. He, N.E. Sharpless, Senescence in Health and Disease, Cell, 169 (2017) 1000–1011.

[49] Y. Zhu, T. Tchkonia, T. Pirtskhalava, A.C. Gower, H. Ding, N. Giorgadze, A.K. Palmer, Y. Ikeno, G.B. Hubbard, M. Lenburg, S.P. O’Hara, N.F. LaRusso, J.D. Miller, C.M. Roos, G.C. Verzosa, N.K. LeBrasseur, J.D. Wren, J.N. Farr, S. Khosla, M.B. Stout, S.J. McGowan, H. Fuhrmann-Stroissnigg, A.U. Gurkar, J. Zhao, D. Colangelo, A. Dorronsoro, Y.Y. Ling, A.S. Barghouthy, D.C. Navarro, T. Sano, P.D. Robbins, L.J. Niedernhofer, J.L. Kirkland, The Achilles’ heel of senescent cells: from transcriptome to senolytic drugs, Aging Cell, 14 (2015) 644–658.

[50] J.D. Leverson, D.C. Phillips, M.J. Mitten, E.R. Boghaert, D. Diaz, S.K. Tahir, L.D. Belmont, P. Nimmer, Y. Xiao, X.M. Ma, K.N. Lowes, P. Kovar, J. Chen, S. Jin, M. Smith, J. Xue, H. Zhang, A. Oleksijew, T.J. Magoc, K.S. Vaidya, D.H. Albert, J.M. Tarrant, N. La, L. Wang, Z.F. Tao, M.D. Wendt, D. Sampath, S.H. Rosenberg, C. Tse, D.C. Huang, W.J. Fairbrother, S.W. Elmore, A.J. Souers, Exploiting selective BCL-2 family inhibitors to dissect cell survival dependencies and define improved strategies for cancer therapy, Sci Transl Med, 7 (2015) 279ra240.

[51] H. Zhu, Y. Tian, H. Chen, Y. Qian, J. Li, D. Niu, W. Zhao, Y. Wu, X. Zhang, T. Tang, H. Li, Y.F. Xian, D. Sun, C. Jin, Targeting DNA damage response pathways in tumor drug resistance: Mechanisms, clinical implications, and future directions, Drug Resist Updat, 83 (2025) 101287.

[52] Z. Xu, M. Zhu, L. Geng, J. Zhang, J. Xia, Q. Wang, H. An, A. Xia, Y. Yu, S. Liu, J. Tong, W.G. Zhu, Y. Jiang, B. Sun, Targeting NAT10 attenuates homologous recombination via destabilizing DNA:RNA hybrids and overcomes PARP inhibitor resistance in cancers, Drug Resist Updat, 81 (2025) 101241.

[53] X. Peng, Y. Wang, Z. Yu, S. Huang, S. Zhang, Z. Zhong, Y. Wang, S. Liu, K. Wang, C. Nicot, F.X. Claret, D. Kong, Jab1 regulates HRR mRNA stability to modulate PARP inhibitor sensitivity in triple-negative breast cancer, Mol Cancer, 24 (2025) 217.

