## Supplementary material for "Targeting Cellular Pseudo-Senescence to Overcome PARP inhibitors Resistance in *BRCA1*-Mutated Breast Cancer": Combined Supplementary Figures

**Supplementary Fig1. Heatmap of SAUL_SEN_MAYO signature gene expression in *ZNF251*WT and KD cells treated with Olaparib.**

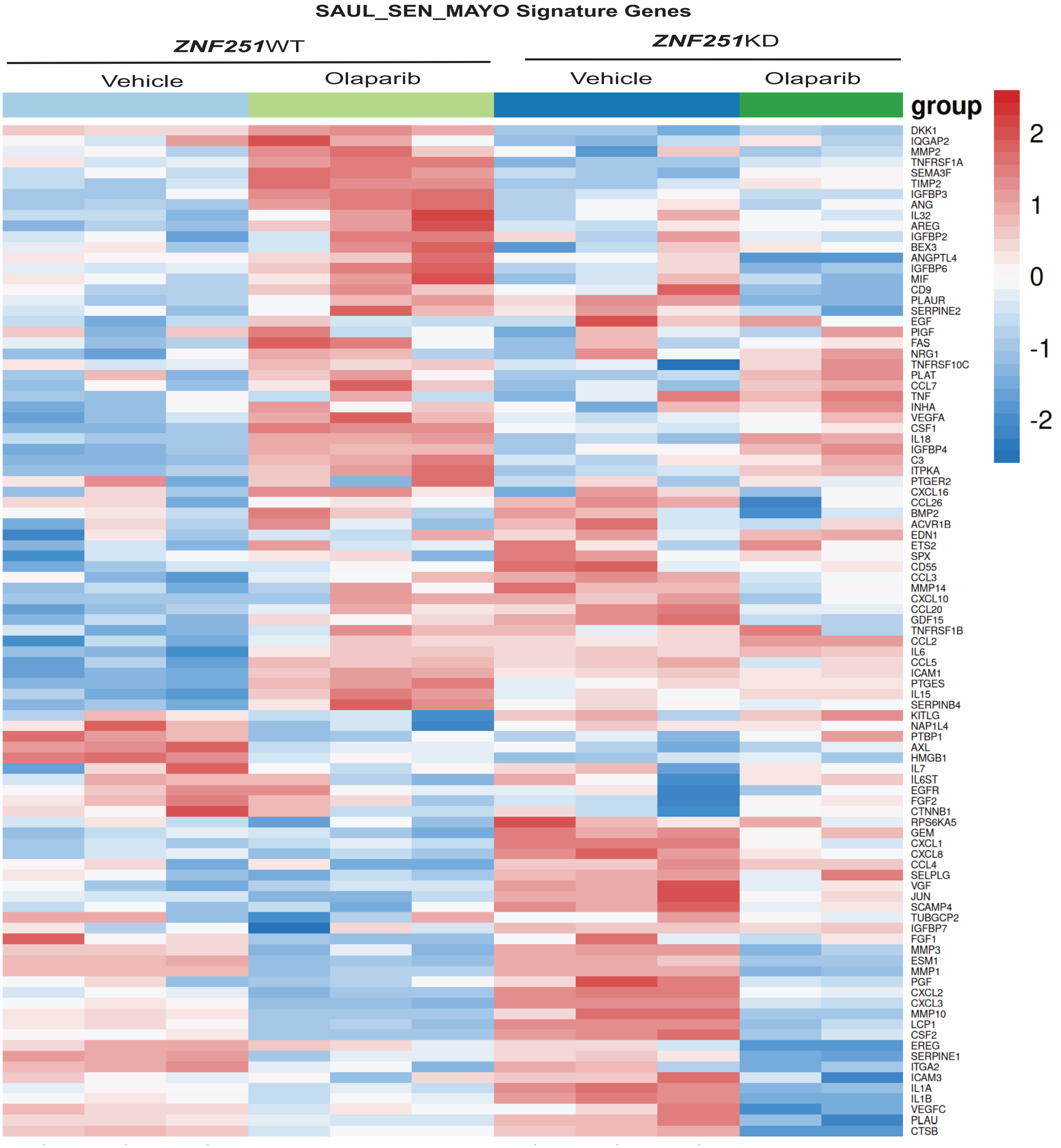


Supplementary Fig1. *ZNF251*KD deficiency induces senescence-associated transcriptional signatures in *BRCA1*-mutant breast cancer cells. Heatmap showing the expression of genes from the SAUL_SEN_MAYO senescence-associated gene signature in *ZNF251*WT and *ZNF251*KD *BRCA1*-mutant breast cancer cells treated with vehicle or olaparib. Gene expression is presented as row-wise Z-scores (red, high; blue, low). *ZNF251*KD cells displayed enrichment of senescence-associated gene expression relative to WT cells under both treatment conditions, supporting induction of a senescence-associated transcriptional state following *ZNF251* loss.

**Supplementary Fig2. Heatmap of GSVA enrichment scores for senescence-related pathways in *ZNF251*WT and KD cells treated with Olaparib.**

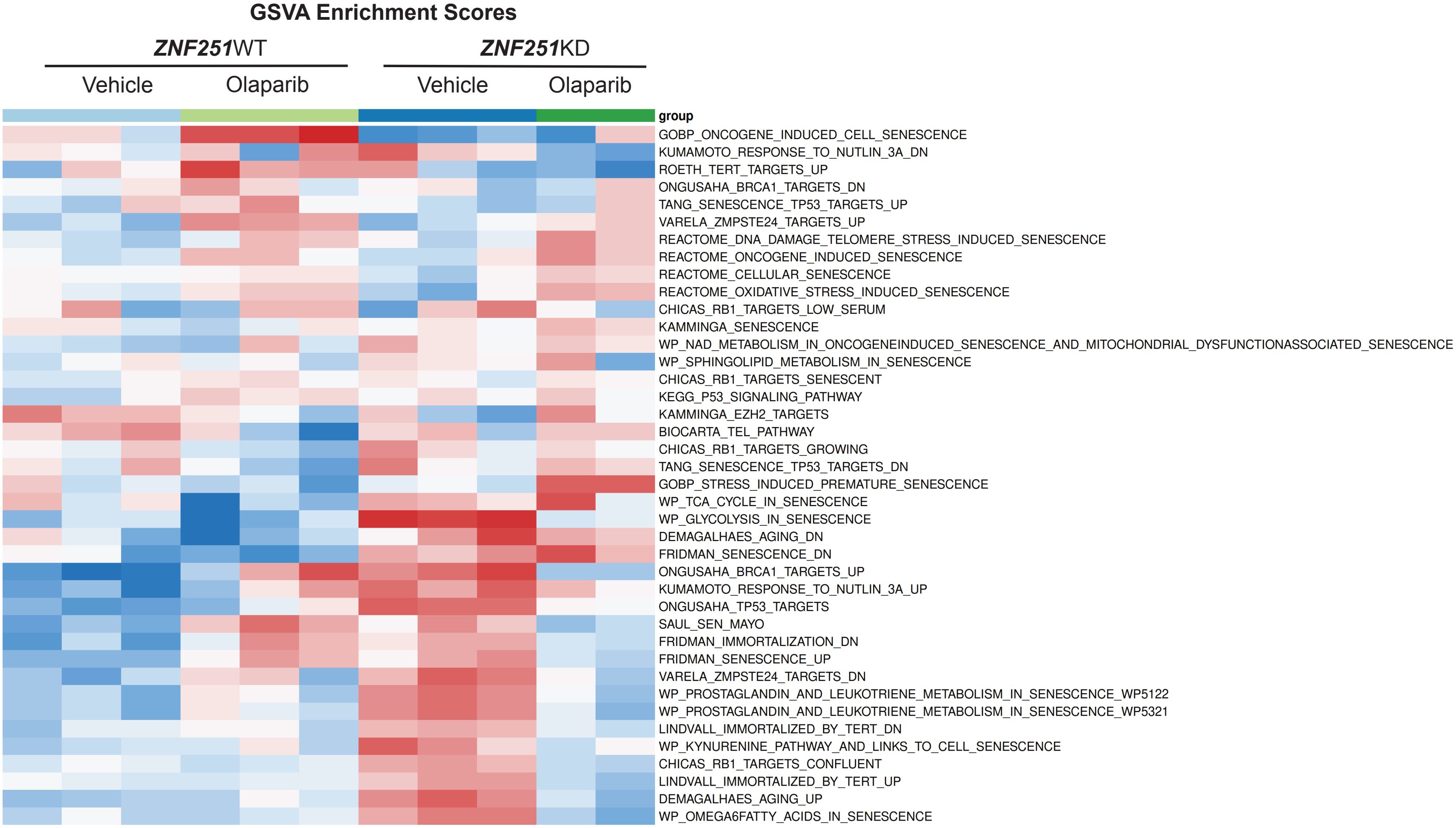


Supplementary Fig2. GSVA analysis demonstrates enrichment of senescence-associated transcriptional programs in *ZNF251*-deficient *BRCA1*-mutant breast cancer cells. Heatmap of GSVA enrichment scores for curated senescence-associated gene sets in *ZNF251*WT and *ZNF251*KD *BRCA1*-mutant breast cancer cells treated with vehicle or olaparib. Enrichment scores were normalized across samples and are shown as row-wise Z-scores (red, higher enrichment; blue, lower enrichment). Multiple independent senescence-related gene signatures were enriched in *ZNF251*KD cells relative to WT cells, supporting induction of a senescence-associated transcriptional state following *ZNF251* loss.

Figure S2

**Supplementary Fig3. *ZNF251*KD amplifies olaparib-induced inflammatory cytokine and p21 expression in MDA-MB-436 and HCC1937 breast cancer cells.**

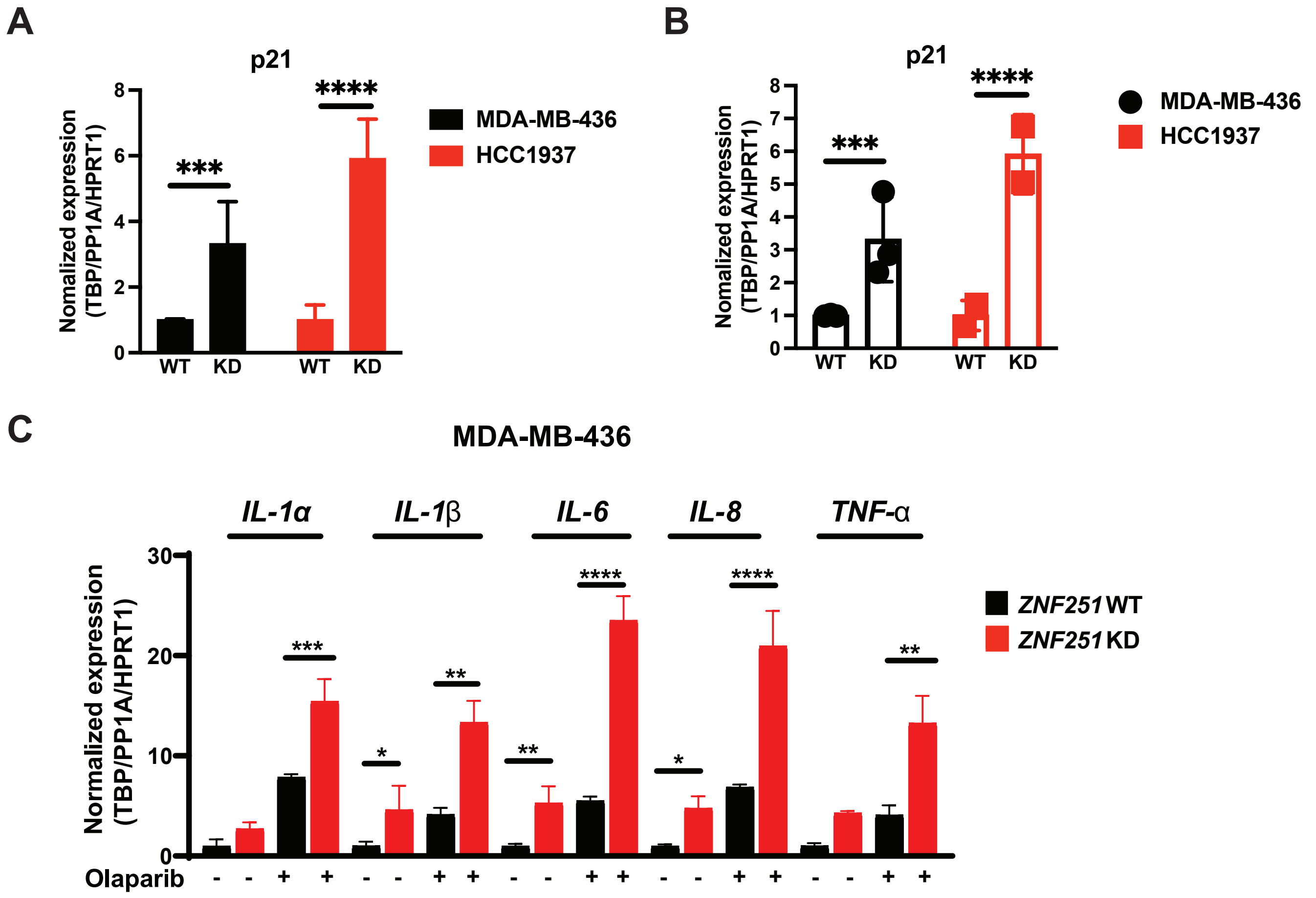


Supplementary Figure 3. A. RT-qPCR of p21 (CDKN1A) mRNA in *ZNF251*WT and KD MDA-MB-436 (black) and HCC1937 (red) cells as bar graph. Expression was normalized to *TBP*, *PPIA*, and *HPRT1* and is shown relative to WT. Each point is one independent experiment (n = 3); bars show mean ± SD, ***P < 0.001, ****P < 0.0001. B. RT-qPCR of p21 (CDKN1A) mRNA in *ZNF251*WT and KD MDA-MB-436 (black) and HCC1937 (red) cells, shown as a scatter dot plot. C. RT-qPCR analysis of *IL-1α, IL-1β, IL-6, IL-8, and TNF-α* mRNA in *ZNF251*WT (black) and *ZNF251*KD (red) MDA-MB-436 cells treated with vehicle (−) or olaparib (+,8uM) for 3 days. Expression was normalized to the geometric mean of TBP, PPIA, and HPRT1 and is shown relative to vehicle-treated WT cells. Data are mean ± SD of n = 3 independent experiments, each run in technical triplicate. Statistical significance was assessed by two-way ANOVA; horizontal bars indicate significant comparisons (*p<0.05, **  p<0.01, *** p<0.001, ****p<0.0001).

**Supplementary Fig4. *ZNF251*KD–induced PARPi resistance in *BRCA1*-mutant breast cancer cells is reversed by NF-κB or BCL-2 family inhibition.**

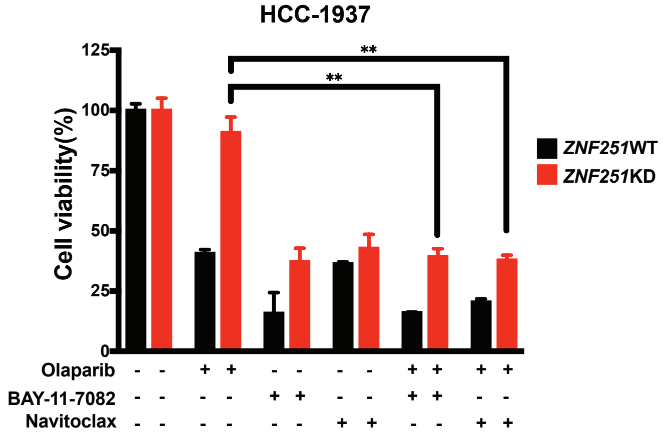


Supplementary Figure 4. NF-κB or BCL-2 family inhibition reverses olaparib resistance in *ZNF251*-depleted HCC1937 cells. *ZNF251*WT and *ZNF251*KD HCC1937 *BRCA1*-mutant breast cancer cells were treated with olaparib, BAY 11-7082, or navitoclax alone, or with the indicated combinations, and cell viability was assessed. Olaparib alone reduced viability in *ZNF251*WT cells but had little effect in *ZNF251*KD cells, consistent with the olaparib-resistant phenotype. Addition of either BAY 11-7082 or navitoclax restored olaparib sensitivity in *ZNF251*KD cells. Data are presented as mean ± SD of three independent experiments. **P < 0.01.

**Supplementary Figure 5. BCL-2 family inhibitors improved the sensitivity to olaparib in breast cancer patient samples.**

A

| Case ID | Age | Gender | Detailed clinical Diagnosis | Staging | Clinical site provided test results |
|---|---|---|---|---|---|
| 131484 | 74 | Female | Invasive ductal Carcinoma | IIA | Her-2/neu: Negative<br>ER: Positive<br>PR: Negative |
| 143814 | 43 | Female | Invasive ductal Carcinoma | IIB | NA |
| 137993 | 57 | Female | Breast adenocarcinoma | IIIC | Her-2/neu: Negative<br>ER: Negative<br>PR: Negative |

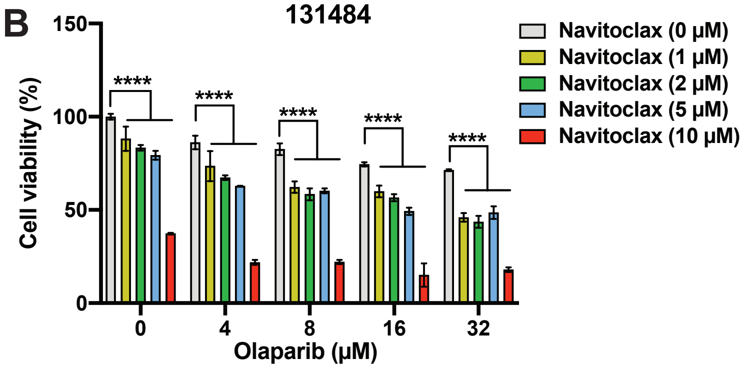


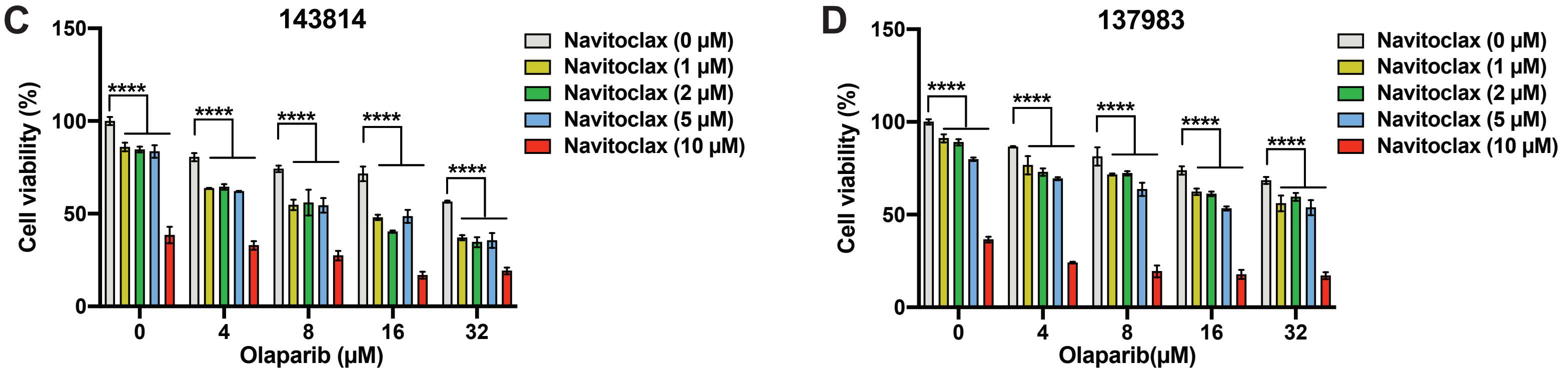


Supplementary Figure 5. (A) Clinical characteristics of three breast cancer patient-derived cell samples used in this study. (B-D) Cell viability analysis of patient-derived breast cancer cells 131484 (B), 143814 (C), and 137983 (D) treated with increasing concentrations of olaparib in combination with navitoclax. Combination treatment with navitoclax and olaparib significantly reduced cell viability compared with olaparib monotherapy, indicating enhanced sensitivity to PARP inhibition. *BRCA* status was not determined by sequencing for any of the three samples.
